# Genetic Screen in a Pre-Clinical Model of High-Grade Complex Karyotype Sarcoma Characterizes Drivers of Distinct Sarcoma Subtypes and Identifies New Therapeutic Vulnerabilities

**DOI:** 10.1101/2022.06.13.495815

**Authors:** Jack Freeland, Maria Muñoz, Edmond O’Donnell, Justin Langerman, Jessica Bergonio, Julissa Suarez-Navarro, Morgan Darrow, Steven Thorpe, Robert Canter, R. Lor Randall, Kathrin Plath, Kermit Carraway, Owen N. Witte, Thomas G. Graeber, Janai R. Carr-Ascher

## Abstract

High-grade complex karyotype sarcomas are a heterogeneous group of more than seventy tumors that vary in histology, clinical course, and patient demographics. Despite these clear differences, these high-grade sarcomas are treated similarly with a uniformly high metastatic rate. Pre-clinical models that allow for rigorous comparisons of distinct human sarcoma subtypes would advance insights into the relationships between sarcomas and inform therapeutic decisions. We describe the robust transformation of human mesenchymal stem cells into multiple subtypes of high-grade sarcoma. Using a pooled genetic screening approach, we identified key drivers and potential modifiers of transformation. *YAP1* and *KRAS* were validated as drivers of two distinct sarcoma subtypes, undifferentiated pleomorphic sarcoma (UPS) and myxofibrosarcoma (MFS), respectively. In addition, the pathology of tumors driven by *CDK4* and *PIK3CA* reflected leiomyosarcoma (LMS) and osteosarcoma (OS) indicating that further iterations of this model could result in additional sarcoma subtypes. Histologically and phenotypically these tumors reflect human sarcomas including the pathognomonic complex karyotype. In addition, *CDK4* and *PIK3CA* driven tumors demonstrated endogenous *YAP1* amplification which is seen across a subset of human tumors. While all tumors overlapped transcriptionally with the TCGA sarcoma data, further analysis confirmed that *YAP1* and *KRAS* tumors recapitulate the UPS and MFS subtypes. Co-analysis of TCGA and model tumors support that these sarcoma subtypes lie along a spectrum of disease and adds guidance for further transcriptome-based refinement of sarcoma subtyping. Within complex karyotype sarcomas, there are multiple genetic changes but identifying those that are clinically relevant has been challenging. Comparing differentially expressed genes in *YAP1* and *KRAS* tumors to human UPS and MFS identified the enrichment of oxidative phosphorylation pathways in both *YAP1* tumors and UPS. Treatment of a panel of sarcoma cell lines with the combination of an oxidative phosphorylation inhibitor and Hippo pathway inhibitor led to a significant impairment in growth identifying new therapeutic targets. A subset of human UPS tumors showed an even greater enrichment in these pathways indicating this model can be used to identify clinically relevant subtypes. This model can be used to begin to understand pathways and mechanisms driving human sarcoma development, the relationship between sarcoma subtypes and to identify and test new therapeutic vulnerabilities for this aggressive and heterogeneous disease.

**Statement of Significance:** We have created the first model to study the development, growth, and metastasis of multiple human sarcoma subtypes. This system can be used as a platform to investigate sarcoma biology and identify new therapeutic targets across a heterogeneous disease.

## Introduction

High-grade complex karyotype sarcomas are a heterogeneous group of tumors with a 65% survival rate at 5 years despite aggressive multimodality treatment with surgery, radiation, and chemotherapy. The heterogeneity of the disease is evidenced by the more than 70 sarcoma subtypes with varying histology, genetics, and patient demographics leading to challenges in treatment and clinical trial design ^1^.

High-grade sarcomas can be genetically subcategorized into two distinct groups. First, those that have a simple karyotype where the biology is driven by a known translocation^2^. Second, complex karyotype sarcomas display an aneuploidy phenotype with numerous chromosomal gains and losses^3^. In adults, 85% of sarcomas are complex karyotype, including undifferentiated pleomorphic sarcoma (UPS), a highly aggressive tumor characterized by a lack of specific markers of differentiation. Contrary to translocation driven sarcomas that have a clearly defined oncogenic driver, pathways and factors leading to the development of complex karyotype sarcomas are poorly understood. As a result, clinically, these complex karyotype sarcomas are treated similarly with varying degrees of success. Further contributing to the complexity of sarcoma management is the rarity of sarcomas as they cumulatively constitute less than 1% of all tumors leading to limited biological material for study and small sample sizes for clinical trials resulting in a generalization of treatments for a heterogeneous family of diseases^4^. The development of pre-clinical models that recapitulate the genetic and biological heterogeneity of individual sarcoma subtypes and allow for the identification of critical genes and pathways driving the pathophysiology of specific sarcoma subtypes would promote discovery of subtype specific and pan-sarcoma therapeutic targets.

To gain insight into the biology of complex karyotype soft tissue sarcomas, The Cancer Genome Atlas (TCGA) genetically characterized the most common sarcomas in adults. These tumors were found to have a low mutation burden and a high rate of aneuploidy^5^. The most common mutations and chromosomal alterations were loss of known tumor suppressors *RB1* or *P53* which have established roles in sarcoma biology ^3,6,7^. Incorporating TCGA data as well as other large datasets, *P53* was found to be altered in 47% of sarcomas. *RB1* mutation or deletion was seen in 22% of tumors with the majority of these also carrying an alteration in P53^8^. These studies indicate that *RB1* and *P53* play important roles in human sarcoma biology. This has also been shown in mouse models where targeted deletion of *p53* drives sarcoma formation^6,9^.

Mesenchymal stem cells (MSCs) give rise to the connective tissues of the body and are the presumed cell of origin for bone and soft tissue sarcomas^10^. In mouse studies, loss of *p53* in mesenchymal stem cells was sufficient to drive sarcoma formation ^6,11^. In contrast, loss of *P53* alone is insufficient to induce sarcoma formation from human mesenchymal stem cells^12^. Prior models of human sarcoma using MSCs as the cell of origin have relied primarily on viral proteins such as SV40 or mutated *RAS* to drive transformation, and these have resulted in the formation of a single undifferentiated sarcoma subtype^13–17^. More recently, human MSCs have been transformed to osteosarcoma using *RB1* silencing and *c-Myc* overexpression or by the addition of *c-JUN* to E6/E7 immortalized MSCs^18,19^. While these models have provided valuable insights, the ability to make comparisons among human sarcoma subtypes within a common model system has not been possible.

Here, we describe the development and characterization of a new pre-clinical model of human sarcoma. This system transforms mesenchymal stem cells into high-grade sarcomas by introducing genetic alterations observed across human sarcomas. Histologically, this results in the formation of four distinct high grade sarcoma subtypes, namely, undifferentiated pleomorphic sarcoma (UPS), myxofibrosarcoma (MFS), osteosarcoma (OS), and leiomyosarcoma (LMS) all of which contain a complex karyotype or aneuploidy phenotype. Further transcriptional analysis demonstrated that UPS and MFS genetically reflect the human disease and comparisons of the model to human tumors identified clinically relevant subpopulations and new therapeutic opportunities.

## Results

### Human Mesenchymal Stem Cells can be Transformed to High-Grade Sarcomas

To develop an i*n vivo* model of sarcoma development with the goal of creating different subtypes, we began with human MSCs and aimed to introduce genetic changes that are commonly observed across high grade sarcomas (Figure 1A). While mouse MSCs can be readily transformed to sarcomas, human MSCs have been more challenging^12^. A putative mechanism for this variation between species is due to differences in telomere maintenance. This is exemplified by the successful transformation of human MSCs when *hTERT* has been added along with viral proteins or mutated *H-RAS* ^15–17^. *hTERT* was demonstrated in the TCGA data to be amplified in a subset of human sarcoma^5^. Therefore, as a basis for these studies, we utilized ASC52telo, a MSC line that has been immortalized by *hTERT* (human telomerase) ^20^. Despite immortalization, these cells have identical cell surface marker expression to non-immortalized control mesenchymal stem cells (Supplemental Figure 1A). These cells can undergo in vitro differentiation into adipocytes, osteocytes, and chondrocytes. Adipogenic differentiation is limited compared to non-immortalized MSCs due to impaired contact inhibition and chondrogenic pellets were larger and more diffuse due to an increased proliferative rate (Supplemental Figure 1B).

**Figure 1.**
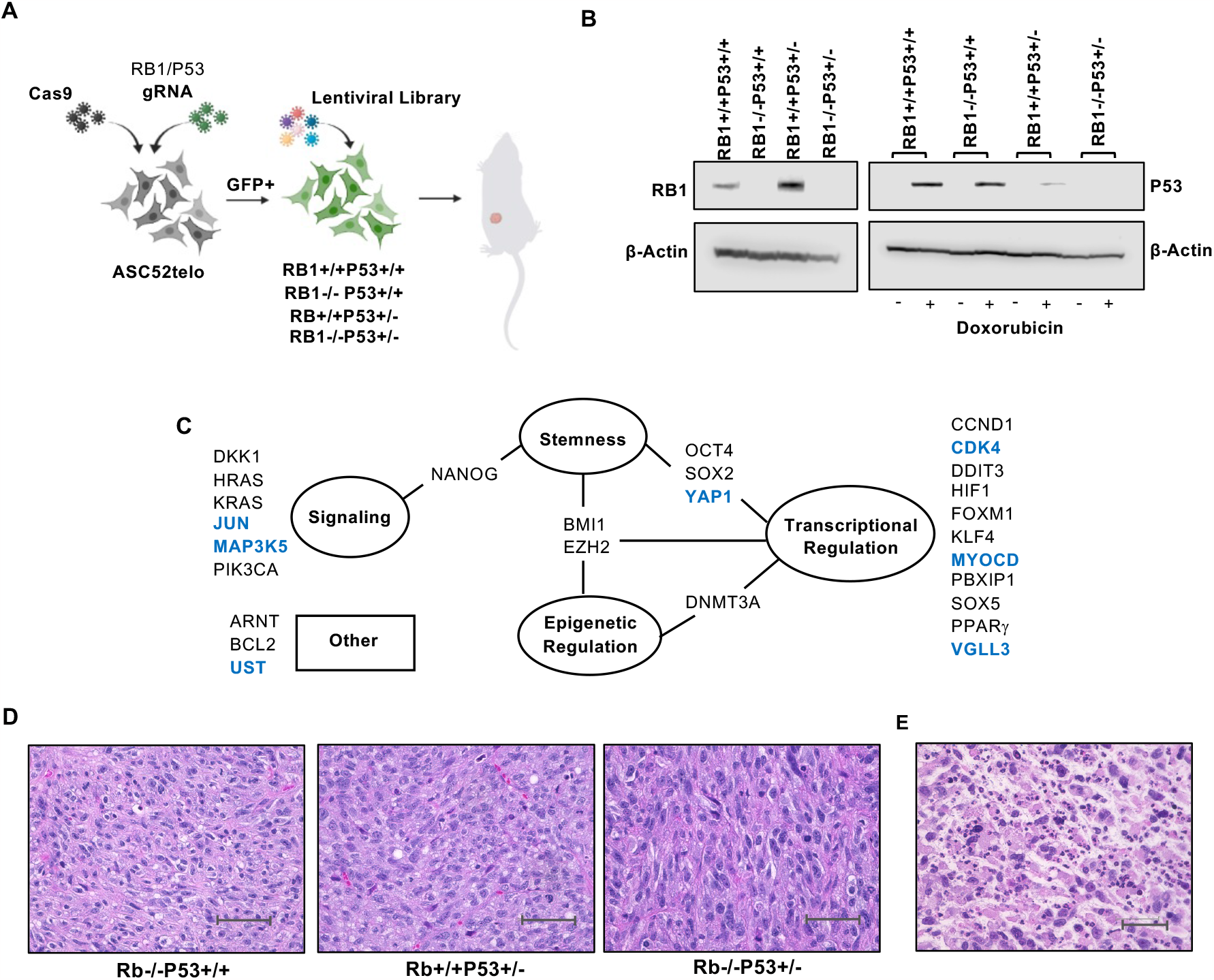
Human Mesenchymal Stem Cells can be Transformed to High-Grade Sarcomas. (A) Schematic diagram of forward transformation model of sarcoma development. (B) Western blot of generated cell lines were evaluated for RB1 and P53 expression. Given the low basal levels of P53, cells were treated with doxorubicin to induce expression. Beta actin is shown as a loading control. Experiment was performed in duplicate (C) Schematic of genes included in the lentiviral library separated into groups based on known functions. (D) High power views of sarcomas formed by addition of the lentiviral vector to each of the genetic backgrounds shown. Scale bar represents 50um. (E) Distinct areas of necrosis were seen as evidenced by an immune infiltrate, scale bar represents 50um. N=8 per genotype.

*RB1* and *P53* are commonly inactivated or mutated in human soft tissue sarcomas^3^. There is also evidence for amplification of negative regulators of these genes, further emphasizing their importance in sarcoma biology^5^. Therefore, we targeted *RB1* and *P53* in ASC52telo using a CRISPR-Cas9 system ^21^. A lentivirus containing doxycycline-inducible Cas9 was transduced into the ASC52telo cell line ^22^. An inducible system was used to mitigate the potential *in vitro* and *in vivo* toxicity of high levels of Cas9 ^23^. Single cell clones were expanded and tested to ensure tight regulation of Cas9 (Supplemental Figure 1C). Guides targeting either *RB1, P53*, both genes, or a control sequence were inserted into a lentiviral vector containing GFP and cells were sorted and expanded ^24^(Supplemental Figure 1D). Gene knockout was verified at the DNA and protein levels (Figure 1B and Supplemental Figure 1E). This resulted in four cell lines; wildtype (*RB1+/+P53+/+)*, loss of *RB1* alone (*RB-/-P53+/+)*, loss of *P53* alone (*RB+/+P53+/-)* and loss of both genes (*RB-/-P53+/-)*. Interestingly, although numerous clones screened showed heterozygous loss of *P53*, we did not observe homozygous allele loss. Despite this, the *P53* pathway is inactive as evidenced by the lack of expression in response to doxorubicin treatment (Figure 1B).

ASC52telo cells with loss of *RB1, P53*, or both genes failed to yield tumors within 90 days after subcutaneous injection into mice, suggesting that targeting of *RB1* and *P53* alone in the background of *hTERT* immortalization is not sufficient to drive robust tumor formation (0/12 tumors formed). We hypothesized that additional genetic alterations were needed for transformation. To aid in the assessment of multiple candidates, we generated a lentiviral library that would allow for a genetic screen approach. All genes were wildtype given the low mutation rate observed in human samples. Genes upregulated or amplified in the TCGA soft tissue sarcoma dataset such as *JUN, YAP1*, and *CDK4* were included^5^. In addition, genes implicated in the development of sarcomas such as the Wnt antagonist *DKK1* that has been shown to transform human MSCs expressing SV40 to sarcoma were included ^14^. Factors such as *PI3K* and *NANOG* that have been shown to regulate the sarcoma stem cell population were also utilized along with genes involved in the growth of sarcomas such as *FOXM1*^25,26^. Barcoded lentiviral constructs containing twenty-seven genes with a variety of functions including regulation of cellular signaling, stem cell biology, transcription, and epigenetic regulation were used in the screen (Figure 1C, Table 1).

**Table 1.**
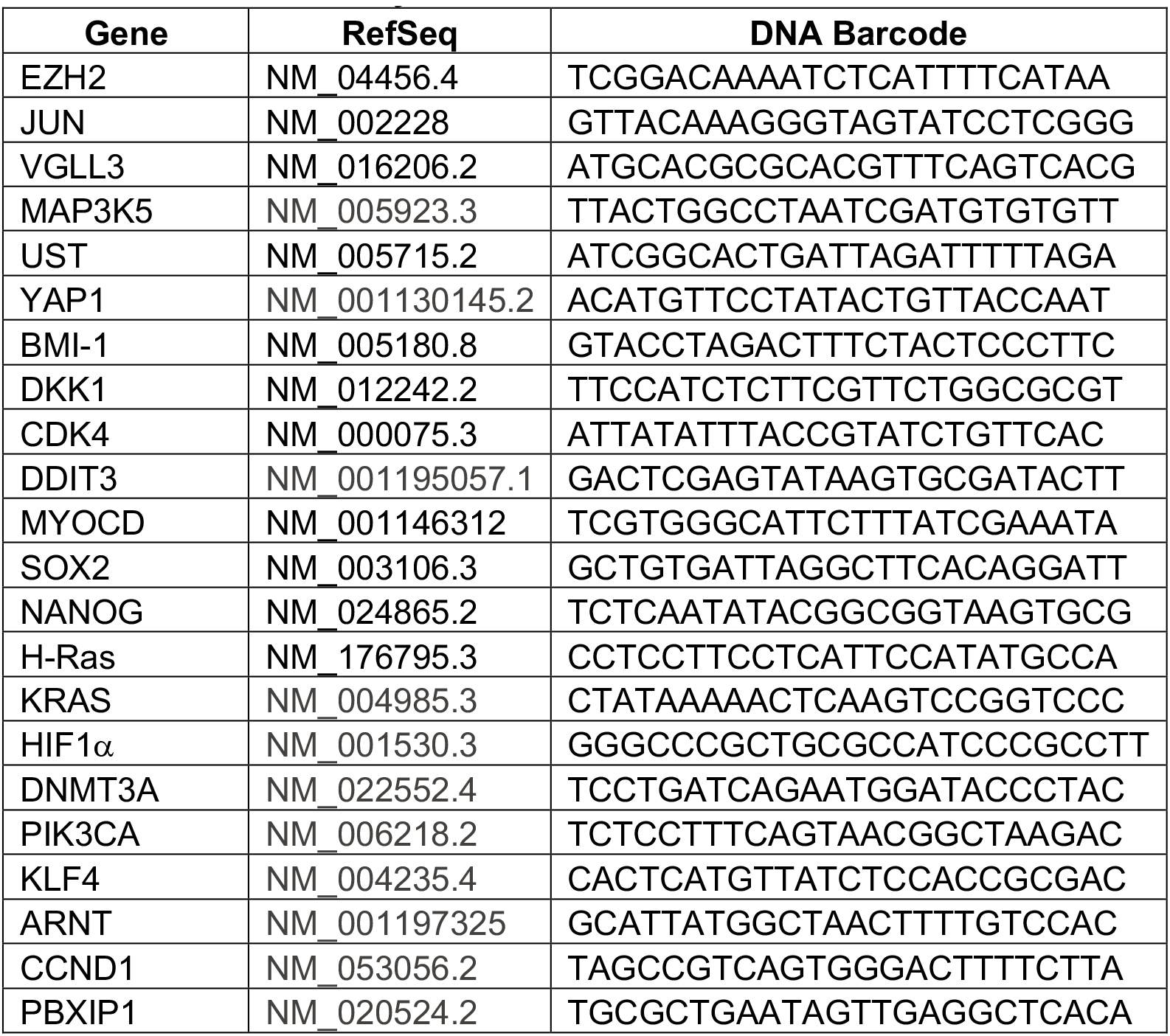
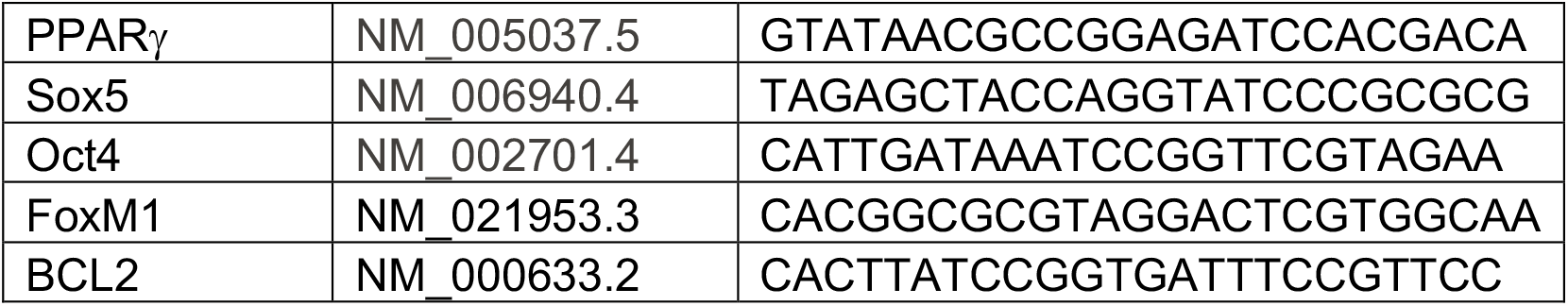
Lentiviral Library.

In murine syngeneic models, loss of *p53* is sufficient for tumor formation but when using human MSCs to induce transformation to high-grade sarcoma, the requirements for *RB1* and *P53* are not known ^27,28^. Therefore, all four lines, *RB1+/+P53+/+, RB1-/-P53+/+, RB1+/+P53+/-* and *RB1-/-P53+/-* cells were transduced with the lentiviral gene library and implanted subcutaneously into immunocompromised non-SCID gamma (NSG) mice. This experiment was performed in duplicate. After 8 weeks, 7/16 (44%) of *RB-/-P53+/+*, 12/16 (75%) of *RB+/+P53+/-* and 14/16 (88%) of *RB1-/-P53+/-* cells transduced with the library formed tumors. Control cells (*RB+/+P53+/+*) did not yield tumors, indicating that the lentiviral library alone is not capable of transforming MSCs to sarcomas. Tumors from these different genetic backgrounds were indistinguishable and demonstrated highly pleomorphic cells with numerous mitotic figures and areas of necrosis consistent with high grade sarcoma (Figure 1D-E). Based on these data, both targeting of a tumor suppressor and activation of an oncogene are needed for transformation of immortalized human mesenchymal stem cells into high-grade sarcomas. For subsequent studies, *RB-/-P53+/-* cells were used given the robust formation of high-grade sarcomas in this background.

### Tumors Formed Histologically and Phenotypically Mimic the Human Disease

In adults, the most common high-grade complex karyotype sarcoma is undifferentiated pleomorphic sarcoma. This tumor type is characterized as having pleomorphic cells and a lack of immunohistochemical staining that would indicate differentiation towards a specific lineage such as bone, muscle, or fat ^1,29^. Further histologic examination of tumors formed from the transduction of the lentiviral library to *RB-/-P53+/-* cells showed primarily areas of high-grade undifferentiated pleomorphic sarcoma although, infrequent areas contained two distinct histologies (Figure 2A). The most abundant histology was consistent with undifferentiated pleomorphic sarcoma (UPS) while the second area contained features of aggressive sarcoma and a lack of differentiation but also abundant myxoid accumulation, consistent with high-grade myxofibrosarcoma (MFS) (Figure 2B-C). Immunohistochemistry of these tumors showed strong positivity for vimentin with alpha smooth muscle actin marking endothelial cells and infrequent tumor cells. Myogenin was consistently negative indicating the tumors did not demonstrate evidence of muscle differentiation that would be consistent with rhabdomyosarcoma (Figure 2D). Taken together, these tumors lack markers of differentiation and represent undifferentiated pleomorphic sarcoma histology with areas of myxofibrosarcoma.

**Figure 2.**
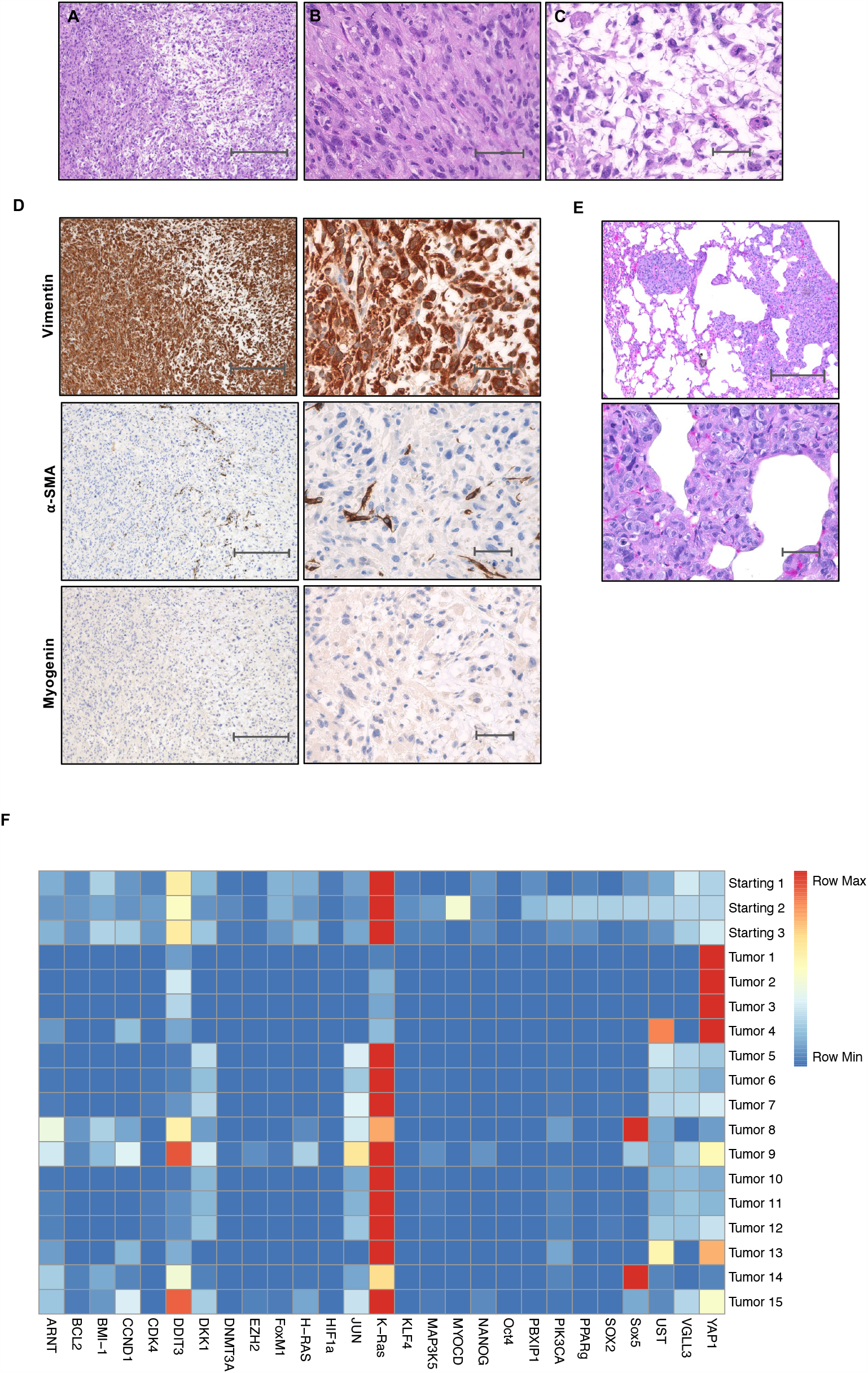
Primary Screen Identifies *YAP1, KRAS*, and *DDIT3* as Potential Drivers of Sarcoma Development. (A) Low power view of H&E staining of tumors formed that demonstrated two distinct histologies. Scale bar represents 300um. High power view of the two distinct histologies demonstrate (B) Undifferentiated pleomorphic sarcoma and (C) Myxofibrosarcoma. Scale bar represents 50uM. Primary screen was performed in triplicate with 4-8 implants per experiment (D) Immunohistochemistry of tumors formed show strong positivity for vimentin, primarily endothelial cell staining with infrequent tumor cells showing positivity for smooth muscle actin, and negative staining for myogenin. Left side shows low power views with scale bars representing 300um and right side shows high power views with 50um scale bars. N=6, two tumors from each experiment were stained for the panel of IHC markers. (E) Representative lung sections show the presence of metastatic sarcoma cells with low (top) and high (bottom) power views and scale bars representing 300um and 50um respectively. N=12 animals total in duplicate experiments, of those analyzed, 5 demonstrated lung metastasis (42%) (F) Heat map representing barcode sequencing counts from the primary screen. See also supplemental figure 2 for percentage of each barcode per sample and calculated p-values. Barcodes were analyzed from distinct tumor samples.

To determine if these tumors phenotypically mimic the human disease, cells were injected into the gastrocnemius muscle of immunocompromised mice. In humans the majority of sarcomas are found in the extremities and metastasize almost exclusively to the lungs ^30^. Consistent with this, implantation of *RB-/-P53+/-* cells with the lentiviral library into the gastrocnemius gave rise to lung metastases in 42% (5/12) of animals without lesions in the lymph nodes, liver, or spleen (Figure 2E). Using this defined and highly efficient system, mesenchymal stem cells can be transformed into high grade sarcomas that histologically and functionally reflect the human disease.

### Primary Screen Identifies *YAP1, KRAS*, and *DDIT3* as Potential Drivers of Sarcoma Development

To identify genes in the lentiviral library that drive high-grade sarcoma formation from MSCs, the introduced genes enriched in the transformation process were identified by sequencing of their associated unique vector barcodes. Of note, all lentiviral library constructs were expressed although at varying reference levels prior to implantation and there was a minimal change in the relative levels over five days of cell culture, indicating significant *in vitro* selection did not occur (Supplemental Figure 2A). Barcode sequencing of the tumor outgrowths from three independent experiments were analyzed and three genes, *YAP1, DDIT3*, and *KRAS* were particularly enriched for and expressed at high levels across multiple tumor samples in three individual experiments (Figure 2F, Supplemental Figure 3A). While *KRAS* and *DDIT3* were strongly represented in the starting lentiviral pool, *YAP1* was not and showed significant enrichment in 13/15 tumors during the process of transformation (Supplemental Figure 3B). This model system allows for the identification of potential drivers of sarcoma development.

### YAP1 and KRAS Drive the Formation of Histologically Distinct Sarcoma Subtypes

*YAP1* or Yes associated protein is a key downstream mediator of Hippo signaling and has been shown to be overexpressed or amplified in sarcomas as well as other cancer types ^25,31,32^. DNA damage inducible transcript 3 (DDIT3) is a member of the C/EBP family of transcription factors and regulates adipogenesis. The *FUS-DDIT3* translocation leads to the formation of myxoid/round cell liposarcoma and amplification of *DDIT3* has been shown to be associated with a myxoid liposarcoma like histology ^33^. Rare mutations in oncogenic *H-RAS* and *KRAS* have been noted in genomic analysis of sarcomas and implicated in transformation, but a role for wildtype *KRAS* has not been established^15,34^. While these factors have roles in sarcoma biology, they have not been experimentally demonstrated to be drivers of human sarcoma development.

Next, a secondary screen was performed to determine if either *YAP1, KRAS*, or *DDIT3* alone were capable of transforming RB-/-P53+/- cells into sarcomas. *RB1-/-P53+/-* cells were transduced with a negative control, blue fluorescent protein (BFP), each lentivirus alone (*YAP1, KRAS*, and *DDIT3*) or the combination as a positive control. These were injected into equal numbers of male and female 6–8-week-old immunocompromised mice. As expected, the combination of all three genes was sufficient to drive transformation. Interestingly, *YAP1* and *KRAS* alone led to tumor formation within 8-16 weeks while addition of *DDIT3* did not after 26 weeks. *YAP1* tumors can be propagated *in vivo* with second passage cells giving rise to tumors within 2 weeks with reproducible histology (Supplemental Figure 2B). Of note, the BFP negative control led to 12.5% implants forming tumors after 26 weeks that histologically represented UPS or osteosarcoma indicating that spontaneous transformation can occur at a long latency.

UPS and MFS exist on a disease spectrum^5^. *YAP1* tumors were histologically consistent with undifferentiated pleomorphic sarcoma as evidenced by vimentin positivity and a lack of expression of markers of differentiated cell types (Figure 3A). *KRAS* tumors had the distinct histological appearance of myxofibrosarcoma (Figure 3B). The latency of tumor development in the secondary screen was longer than that observed for the primary screen indicating that cooperation amongst genes can likely accelerate sarcoma development. In support of this, while *DDIT3* alone did not give rise to tumors within 26 weeks, the addition of *DDIT3* to KRAS tumors led to a more aggressive phenotype with increased pleomorphism and shorter latency (Supplemental Figure 2C) indicating that *DDIT3* is likely a modifier or enhancer of sarcoma development. These data demonstrate that histologically *YAP1* and *KRAS* drive formation of tumors with undifferentiated pleomorphic sarcoma and myxofibrosarcoma histology, respectively.

**Figure 3.**
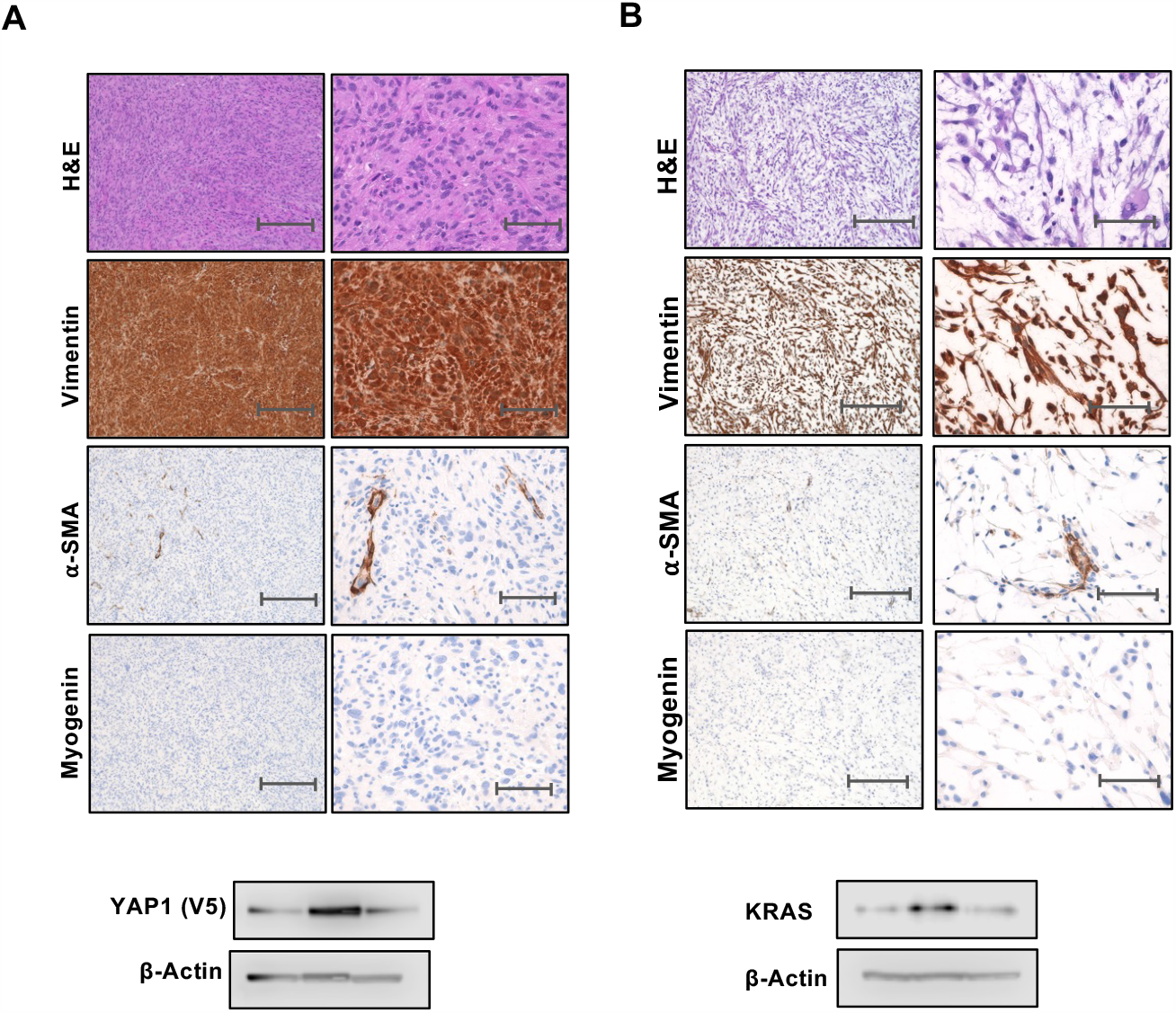
*YAP1* and *KRAS* Drive the Formation of Histologically Distinct Sarcoma Subtypes. (A) Histology from *YAP1* driven tumors are shown with lower power on the left column, scale bars 300um and high power on the right, scale bars 50um. Tumors histologically represent undifferentiated pleomorphic sarcoma with strong staining for vimentin and negative for SMA and myogenin. (B) *KRAS* driven tumors are shown in this panel with histology consistent with myxofibrosarcoma. Low power images are shown on the left with high power on the right. Representative western blots of derived tumors are shown. Secondary screen was performed in duplicate with n=8 for the first experiment and n=4 for the replicate.

### Identification of Additional Drivers and Modifiers of High-Grade Sarcoma Formation

We next considered that additional factors in the lentiviral library could lead to transformation, but this was masked by a proliferative advantage conferred by *KRAS, YAP1*, or *DDIT3*. Therefore, the primary screen was repeated removing *KRAS, YAP1*, and *DDIT3* from the lentiviral library. Histologically, the resulting tumors were most consistent with undifferentiated pleomorphic sarcoma (Figure 4A). Barcode sequencing of these tumors identified several potential drivers. Genes whose barcodes comprised >10% of represented barcodes in one tumor or >5% in two outgrowths were tested individually in a secondary screen. This included *BCL2, CDK4, CCND1, JUN, PIK3CA*, and *SOX2* (Figure 4B).

**Figure 4.**
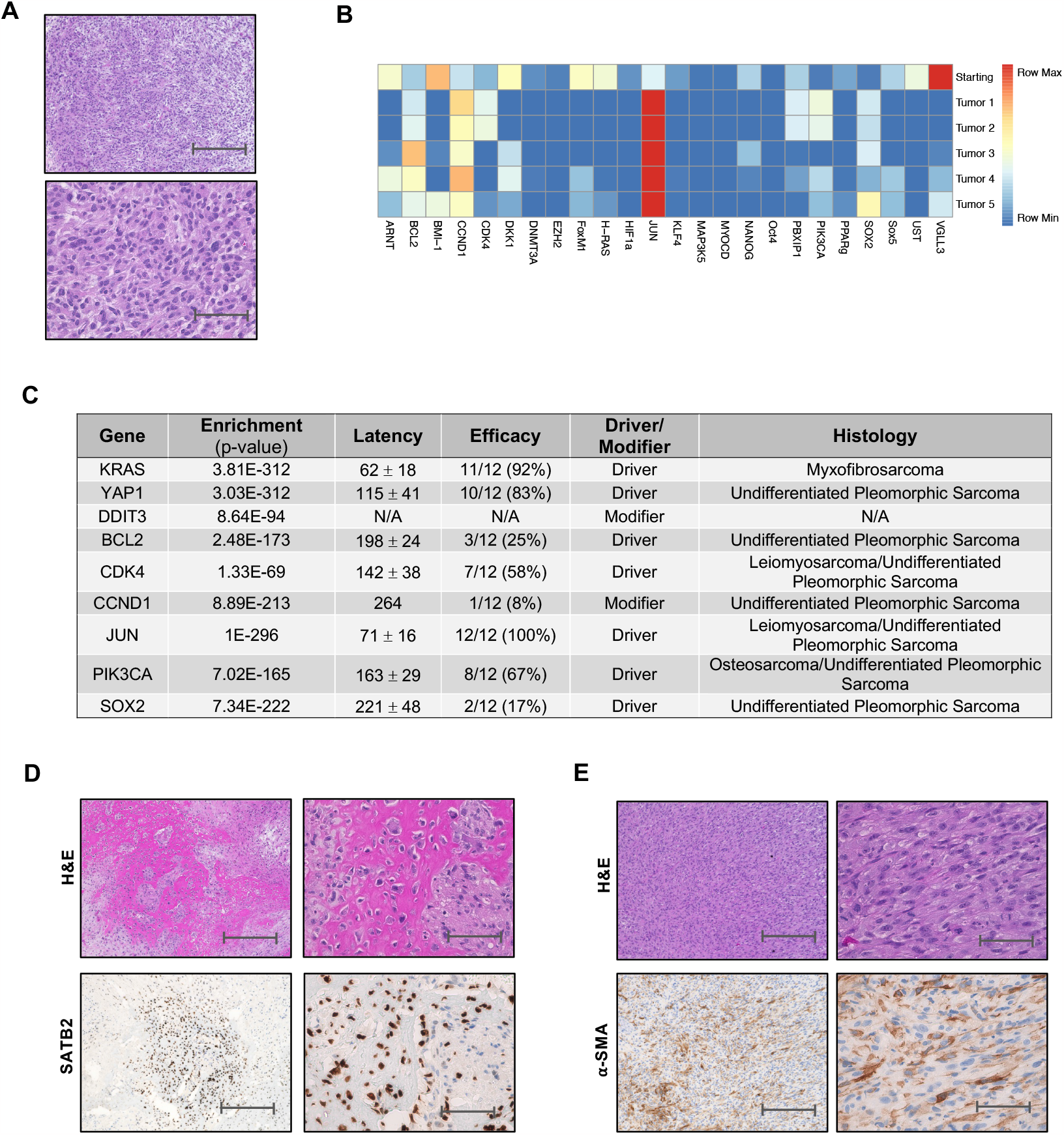
Identification of Additional Drivers of High-Grade Sarcoma Formation. (A) H&E staining of tumors formed from RB1-/-P53+/- cells transduced with the lentiviral library without *YAP1, KRAS*, and *DDIT3*. Low power is shown in the top panel and high power in the lower panel. Experiment was performed in duplicate. (B) Heat map of barcode sequencing from tumor outgrowths. (C) Secondary screen summary table showing the gene that was added to *RB1-/-P53+/-* cells, the amount of time until the tumor reached 1cm (latency) and the number of tumors that formed from the injections (efficacy) as well as the histology of the outgrowths. P-values representing the relative enrichment is shown in the table, for KRAS, YAP1, and DDIT3. Latency is shown as the mean +/- the standard deviation (D) H&E and immunohistochemistry of *PI3KCA* driven tumors that show osteosarcoma confirmed with SATB2 expression. Low power is shown on the left and higher power on the right. (E) H&E and immunohistochemistry of *CDK4* driven tumors showing leiomyosarcoma with positive alpha smooth muscle actin staining. For all low power images, scale bars represent 300um and higher power shows scale bars representing 50um.

*BCL2* and *SOX2* transduced individually gave rise to undifferentiated pleomorphic sarcoma albeit with a significantly prolonged latency and low penetrance as compared to *YAP1* and *KRAS* (Figure 4C). *CCND1* gave rise to one tumor after 264 days and zero tumors in a second experiment after 272 days indicating either a low penetrance or more likely, that the single tumor was a spontaneous transformant and that *CCND1*, like *DDIT3*, is a modifier of sarcoma development and not a strong primary driver.

*JUN* robustly drove the formation of high-grade sarcoma with an average latency of 71 days and 100% penetrance. Amplification or overexpression of *JUN* has been shown to inhibit key drivers of adipocytic differentiation in high-grade sarcoma^35^. Of note, these tumors at <1cm in size were primarily necrotic. Examined sections of viable cells had uniform V5 expression but a highly heterogenous appearance with areas histologically demonstrating undifferentiated pleomorphic sarcoma as well as pleomorphic leiomyosarcoma as evidenced by positive SMA staining and a more spindled morphology (Supplemental Figure 2C).

*PIK3CA* encodes the catalytic subunit of PIK3 which activates Akt signaling. Rarely, mutations of *PIK3CA* have been noted in soft tissue sarcomas and these are associated with poor survival ^36^. Addition of *PI3KCA* to *RB1-/- P53+/-* cells led to formation of tumors although with a lower efficacy and longer latency as compared to cells expressing *YAP1* or *KRAS* (Figure 4C). The histology of 50% of these tumors was consistent with osteosarcoma, confirmed by strong staining for SATB2 (Figure 4D), while 50% represented UPS.

Cyclin dependent kinase 4 (*CDK4*) binds in a complex with cyclin D to phosphorylate RB1 and alleviate repression of E2F to promote cell cycle progression^37^. Alterations in this pathway have been previously noted in large sarcoma sequencing datasets^38^. In TCGA data, CDK4 was shown to be amplified in up to 18.5% of samples^5^. These tumors had a significantly longer latency and a decreased penetrance. Of those, 60% of tumors displayed strong staining of SMA and had a more spindled appearance consistent with pleomorphic leiomyosarcoma while the remaining tumors were histologically UPS (Figure 4E). While the histology of these tumors is not uniform, it does demonstrate the potential of this model and that additional sarcoma subtypes can be generated using this approach.

### Tumor Models Demonstrate an Aneuploidy Phenotype and *YAP1* Amplification

We then sought to determine if these tumors formed genetically reflect the human disease and contain a complex karyotype or aneuploidy phenotype that is pathognomonic of these histologies clinically^1^. To investigate this, *YAP1, KRAS, CDK4*, and *PIK3CA* driven tumors were evaluated by whole exome sequencing with ASC52telo and *RB1-/-P53+/-* pre-transformation parent cells as controls. *JUN* was not included given the histologies were mixed within individual tumors and most tumors were necrotic. Copy number variation (CNV) analysis revealed ASC52telo cells harbor a notable amount of aneuploidy with CNV regions ranging in size from single genes to megabase pairs (Figure 5A). CNV changes are not uncommon in in vitro maintained stem cell cultures, but generally tend to be less than that seen in aneuploid tumors or models^39^. In addition to the CNVs present in ASC52telo cells, which are highly conserved in the tumors, there were tumor model-specific CNVs. For example, *YAP1* driven tumors displayed single-copy number loss of portions of chr 4, 5, 19, and 21 (Supplemental Figure 4A). The *KRAS* driven model displayed single-copy number loss of chr 5 and a portion of chr 19 (Supplemental Figure 4B). The *PIK3CA* driven model displayed single-copy number loss of chr 15 and various smaller deletions (Supplemental Figure 4C). Lastly, the *CDK4* driven model displayed a two-copy number gain of a portion of chr1 where parent *RB1+/+P53+/+* cells displayed a single-copy number gain, indicating that regions of the genome which already contain CNVs in the *RB1+/+P53+/+* cells remain dynamic (Supplemental Figure 4D).

**Figure 5.**
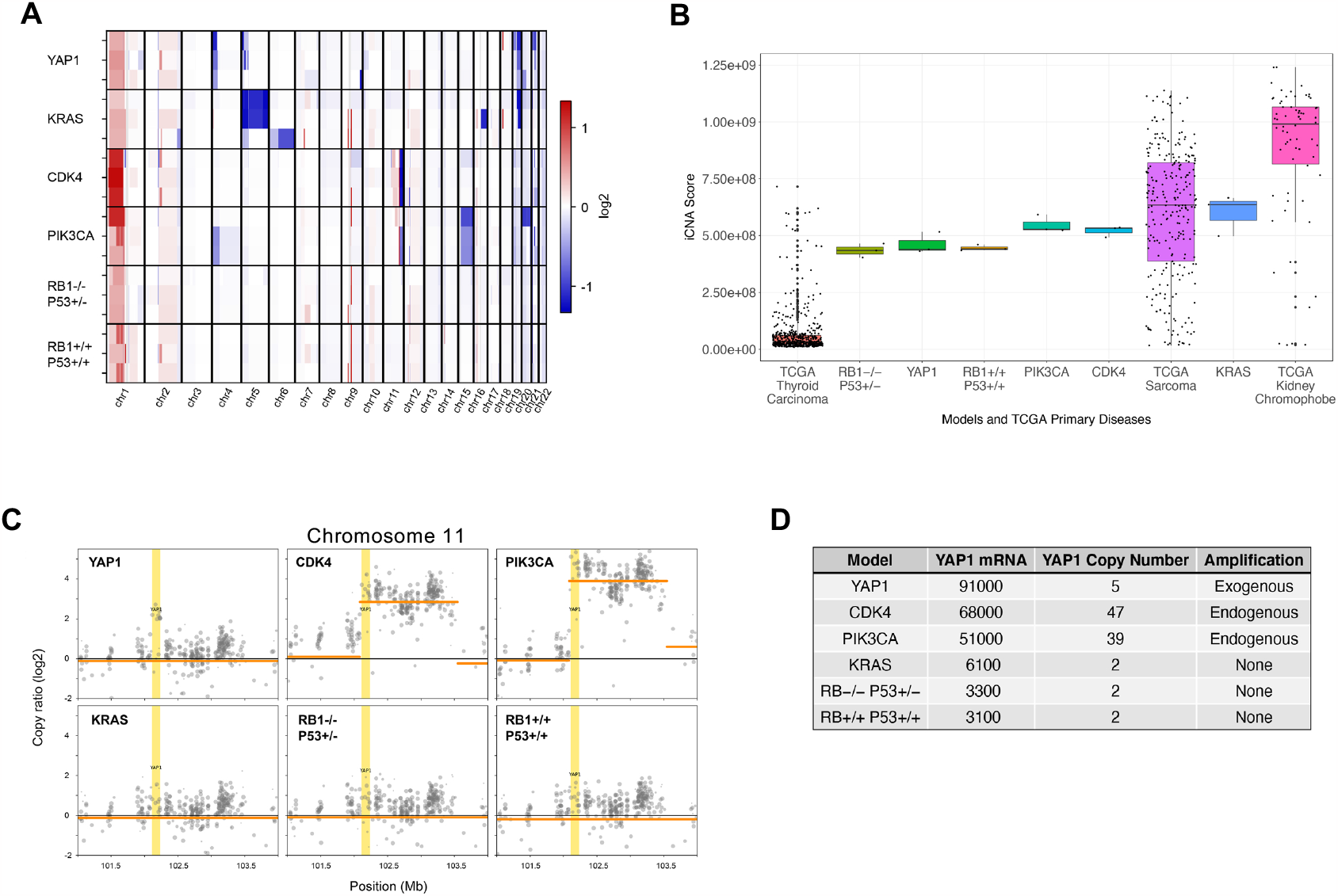
Tumor Models Demonstrate an Aneuploidy Phenotype and *YAP1* Amplification. (A) Heat map showing chromosomal gains (red) and losses (blue) in tumor models and cell lines. Three samples are shown from each tumor type and chromosomes are represented on the X axis. (B) Quantification of aneuploidy by iCNA scores. Thyroid carcinoma and kidney chromophobe are shown on the axis to demonstrate the extremes of the TCGA data. (C) Graph demonstrating chromosome 11q amplification in *CDK4* and *PIK3CA* driven tumors and exogenous amplification of YAP1 in the YAP1 driven tumors. (D) Table summarizing the mRNA expression levels of *YAP1* across the models and the relative copy number.

To quantify the degree of aneuploidy, an integrated copy number alteration (iCNA) score was calculated ^40^. To serve as a relative scale, iCNA scores were also calculated for all 33 TCGA primary tumors (Supplemental Figure 5A). The TCGA data were then ranked by median iCNA score, with the highest and lowest ranking disease, along with sarcoma, being plotted alongside the models and controls (Figure 5B). The tumor model iCNA scores fall within the range of human sarcomas. From these data we can conclude that there is evolution of aneuploidy during the process of transformation and this is consistent with copy number changes contributing to oncogenic fitness and with the predominance of CNVs in the corresponding patient tumors.

*YAP1* has been shown to have copy number gains or to be amplified across a subset of high grade sarcomas^5^. In this pre-clinical model, amplification of chromosome 11q in a 1.5 Mbp region which contains the *YAP1* oncogene was observed (Figure 5C). This amplification was noted across *CDK4* and *PIK3CA* driven tumors with approximately 40-50 copies of the gene present (Figure 5D). Notably, *KRAS* driven tumors did not show amplification or overexpression of *YAP1* as compared to cell lines indicating that the development of these tumors is mechanistically distinct (Figure 5C). Models with endogenous (*CDK4* and *PIK3CA* driven) and exogenous (*YAP1* driven) amplification of *YAP1* show similar *YAP1* gene expression values, while the controls and *KRAS* driven samples are much lower (Figure 5D). It is possible that the YAP1 amplification occurred de novo during transformation or was present as a rare clone in the pre-transformation cells. Nevertheless, this selection for YAP1 signaling indicates it is a key tumor fitness-conferring pathway in sarcoma development and growth.

### Transcriptome of *YAP1* and *KRAS* Driven Tumors Overlaps Human Undifferentiated Pleomorphic Sarcoma and Myxofibrosarcoma

To investigate the transcriptome of the tumors formed in our models and make comparisons to human sarcoma samples, RNA sequencing was performed. In an unsupervised, pan-cancer approach, principal component analysis (PCA) was performed on gene expression data of all 33 TCGA primary diseases (Supplemental Figure 5B). To unbiasedly select a subset of TCGA cancers for comparison, a Euclidean distance in the PC space was calculated to rank each cancer type by overall expression-based closeness or likeness to sarcoma (Table 2). To achieve a range of diseases, every fifth disease plus sarcoma was selected for further analyses. We performed PCA on the TCGA subset and projected the tumors and controls onto the PCA-defined gene expression space (Figure 6A). We saw that both the model tumors and pre-transformed control cells have RNA profiles that are close to sarcoma, indicating the transcriptome of the model reflects that of patient tumors. In addition to sarcoma, our system also overlapped with skin cutaneous melanoma which had ranked number one in distance to sarcoma in the pan-cancer analysis (Table 2). To investigate if the model system preferentially aligned with one of the two cancer types, the model tumors were projected onto a PCA space defined only by skin cutaneous melanoma and sarcoma (Supplemental Figure 5C). Our tumor models projected closer to sarcoma, suggesting their transcriptomes reflect tumors of sarcoma patients (Supplemental Figure 5B)

**Table 2.**

| **Primary Disease** | **Distance** |
| --- | --- |
| Skin cutaneous melanoma | 4.47796483 |
| Adrenocortical cancer | 5.4388277 |
| Mesothelioma | 5.46433186 |
| Thymoma | 5.95929823 |
| Uterine carcinosarcoma | 6.44985246 |
| Testicular germ cell tumor | 6.76786047 |
| Uveal melanoma | 7.64796829 |
| Thyroid carcinoma | 8.11073693 |
| Kidney chromophobe | 8.35292074 |
| Prostate adenocarcinoma | 9.7059719 |
| Uterine corpus endometrioid carcinoma | 9.72997644 |
| Breast invasive carcinoma | 9.79499055 |
| Diffuse large B-cell lymphoma | 9.8110272 |
| Ovarian serous cystadenocarcinoma | 10.3582273 |
| Kidney papillary cell carcinoma | 10.3877392 |
| Kidney clear cell carcinoma | 10.4136214 |
| Acute myeloid leukemia | 12.4056921 |
| Lung adenocarcinoma | 13.0394881 |
| Cholangiocarcinoma | 13.0676807 |
| Pancreatic adenocarcinoma | 13.1890538 |
| Pheochromocytoma & paraganglioma | 13.8051024 |
| Bladder urothelial carcinoma | 14.7929153 |
| Lung squamous cell carcinoma | 14.8189848 |
| Stomach adenocarcinoma | 14.9858603 |
| Glioblastoma multiforme | 15.4837114 |
| Rectum adenocarcinoma | 16.0749137 |
| Colon adenocarcinoma | 16.5443637 |
| Esophageal carcinoma | 16.7102555 |
| Liver hepatocellular carcinoma | 17.4924775 |
| Cervical & endocervical cancer | 18.1199321 |
| Head & neck squamous cell carcinoma | 18.9186762 |
| Brain lower grade glioma | 22.568799 |

**Figure 6.**
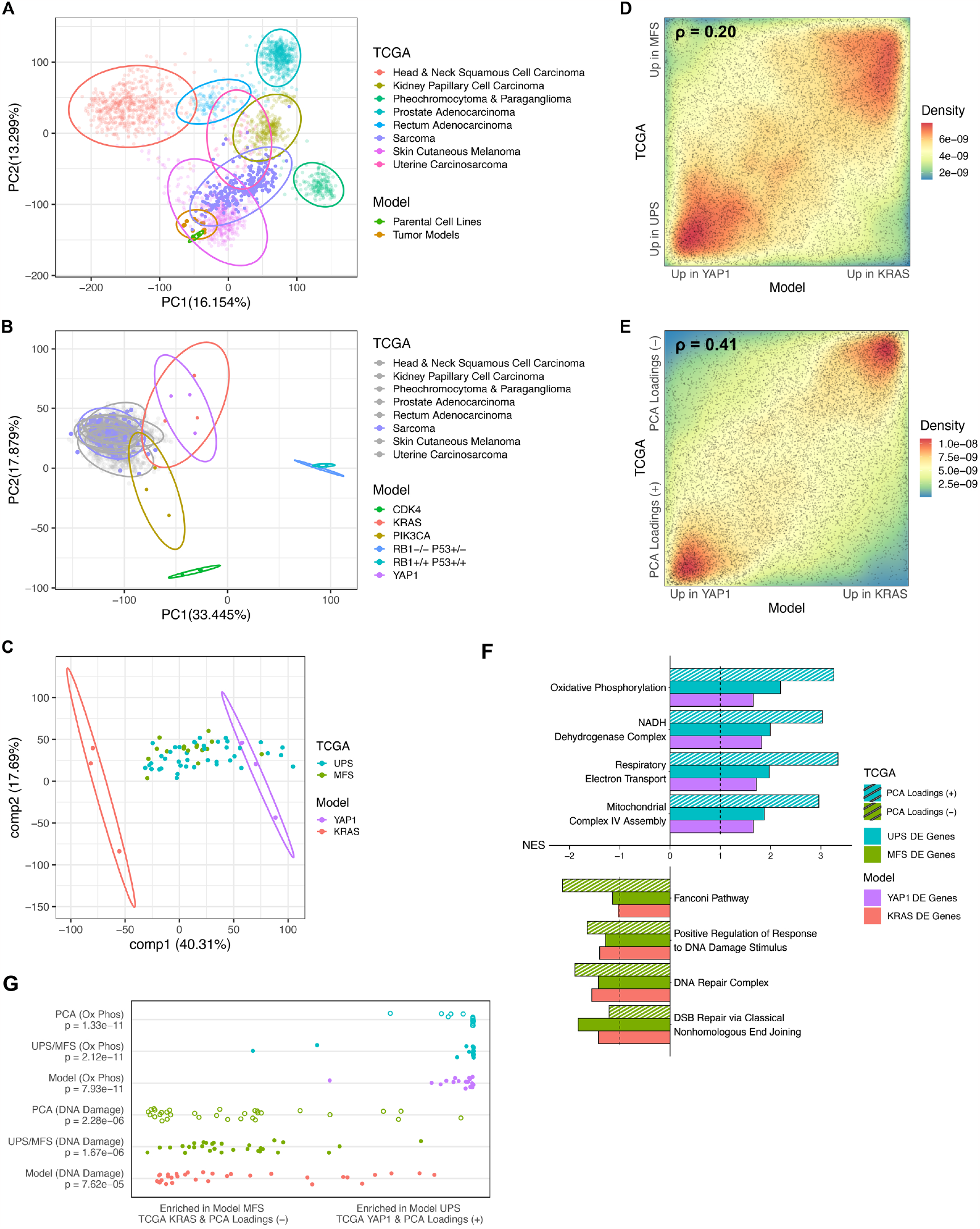
*YAP1* and *KRAS* Driven Tumors Transcriptionally Reflect Human Undifferentiated Pleomorphic Sarcoma and Myxofibrosarcoma. (A) PCA plot of human cancers from TCGA data with projection of the tumor models and cell lines onto the space showing overlap with human sarcomas. (B) PCA plot of tumor models and cell lines with projection of TCGA samples into the space demonstrating that the models cluster with tumors and are distinct from the cell lines. (C) PLSR plot of UPS and MFS transcriptome with *KRAS* and *YAP1* models projected. (D) Co-rank plot showing overlapping signal in KRAS-YAP1 tumor model differentially expressed (DE) gene signature and TCGA MFS-UPS DE gene signature with Pearson correlation coefficient. (E) Co-rank plot showing overlapping signal in KRAS-YAP1 tumor model DE gene signature and TCGA MFS-UPS PCA signature with Pearson correlation coefficient. (F) GSEA comparing YAP1 tumors, human UPS, and PCA loadings (+) to KRAS tumors, human MFS and PCA loadings (-). (G) Distribution of oxidative phosphorylation and DNA damage related pathways in gene sets ranked by normalized enrichment score (NES). All listed categories are nominally significant (p<0.001) by Kolmogorov-Smirnov (KS) test.

As there was a significant overlap between the models and controls in the pan-cancer projection, we investigated to what degree the model tumors were transcriptionally unique and reflected a disease state. PCA performed on the models and pre-transformed control cells revealed a significant difference between the two groups (Figure 6B). Projection of the TCGA tumor subset onto this space resulted in a tight cluster on the tumor model side. This result supports that the separation between tumors and controls is due to the process of transformation and that these models transcriptionally reflect a disease state as compared to controls. It was also observed that while *YAP1* and *KRAS* driven tumors co-occupy a space, *CDK4* and *PIK3CA* driven tumors are transcriptionally distinct.

Further analyses were performed to characterize the *CDK4* and *PIK3CA* driven models. At the given time, we are unable to conclude that these models preferentially reflect leiomyosarcoma and osteosarcoma patient tumors respectively, but simply, sarcoma as a whole. While these tumors were histologically distinct, it is possible that the areas sequenced may contain a mixed histology or these drivers histologically recreate the human disease while the transcriptome is not fully concordant. Gene expression signatures can guide future genetic and other adjustments to the model to improve recapitulation of core human tumor features.

Tumors driven by *YAP1* and *KRAS* consistently demonstrate undifferentiated pleomorphic sarcoma (UPS) and myxofibrosarcoma (MFS). Morphologically, patient tumors from these subtypes are often difficult to distinguish from one another with both tumor types staining negative for markers of differentiation by immunohistochemistry. Previous efforts to molecularly differentiate UPS and MFS have been unsuccessful, suggesting these two phenotypes occupy a disease spectrum rather than two separate entities^5^. To investigate if this spectrum is reflected in our models, MFS and UPS patient data were projected onto a partial least squared regression (PLSR) plot of the *YAP1* and *KRAS* driven models. Qualitatively, the transcriptomes of the two models appear to sit at either endpoint of a UPS/MFS disease spectrum (Figure 6C). In support of the model’s capability to define a translationally relevant spectrum, patient samples exhibited similar distributions in both the model-defined PLSR space and in an unsupervised patient-defined PCA space (Supplemental Figure 6A-B). We additionally observed a subset of UPS samples that do not overlap with the MFS samples, termed “UPS-high,” in both the model and patient-defined spaces. Of note, the UPS-high samples projected closely to our *YAP1* model, further supporting these models as end points of the disease spectrum.

Differential gene expression analysis was performed to further investigate the overlap between models and patient samples. A co-rank plot showing differential gene expression between UPS and MFS patients and between the *YAP1* and *KRAS* driven models identified a statically significant overlap in the patient UPS-MFS and model YAP1-KRAS based signatures. Genes highly expressed in the *YAP1*-driven model were enriched for genes highly expressed in UPS patient tumors, and a corresponding enrichment was observed for the *KRAS-* driven model and MFS patient tumors (Figure 6D). Signature overlap analysis of the co-rank plot using rank-rank hypergeometric overlap (RRHO) yielded a statistically significant signal, supporting that *YAP1* and *KRAS* driven models closely reflect UPS and MFS patients, respectively (Supplemental Figure 6C)^41^.

Further motivated by the challenge of distinguishing MFS and UPS samples histologically, we investigated if gene loadings from the patient-defined PCA space could provide an additional, unbiased and transcriptome-based signature to stratify samples. A co-rank plot between patient-defined PC1 gene loadings and the model YAP1 versus KRAS-based signature resulted in an even higher degree of statistical overlap (Figure 6E). RRHO analysis yielded a strong signal, approximately three times the significance of the histology-based UPS versus MFS signature (Supplemental Figure 6D). Data from both the unbiased and model-based approach supports a spectrum-based approach to classifying tumors in the UPS and MFS subtypes.

In summary, the *YAP1* and *KRAS* model tumors are clearly distinct from the starting untransformed cells and represent a transformed state with complex karyotypes and transcriptomes that recapitulate key aspects of the human disease state of these two distinct sarcoma subtypes.

### Comparison of Model Transcriptome to TCGA Identified New Therapeutic Vulnerabilities

Gene set enrichment analysis (GSEA) revealed the *YAP1* driven model and UPS patient tumors both showed significant enrichment of oxidative phosphorylation related pathways while the KRAS and MFS patient tumors showed enrichment for DNA damage response related pathways (Figure 6F and 6G). Similar enrichment was also observed in the PCA loadings signature. Further examination of the genes contributing to the oxidative phosphorylation pathway enrichment showed upregulation of multiple members of the NDUF (NADPH:Ubiquinone Oxioreductase) family which make up complex I of the electron transport chain. To determine if targeting of oxidative phosphorylation or *YAP* would was clinically relevant, we treated a panel of high-grade soft tissue sarcomas with the Hippo signaling inhibitor Verteporfin or an oxidative phosphorylation inhibitor, IM156. Treatment of cell lines with either drug led to decreased proliferation (Figure 7A and Supplemental Figure 7A and 7B). We then asked if inhibition of both *YAP1* and oxidative phosphorylation would have a synergistic effect in sarcomas. The drug concentrations tested were based on the average IC50 value across the cell lines. As seen in Figure 7B, treatment of cell lines with the combination of IM156 and verteporfin led to a synergistic effect in four out of five lines. This data indicates that targeting of Hippo signaling and oxidative phosphorylation inhibits the growth of a subset of sarcomas and should be further investigated.

**Figure 7.**
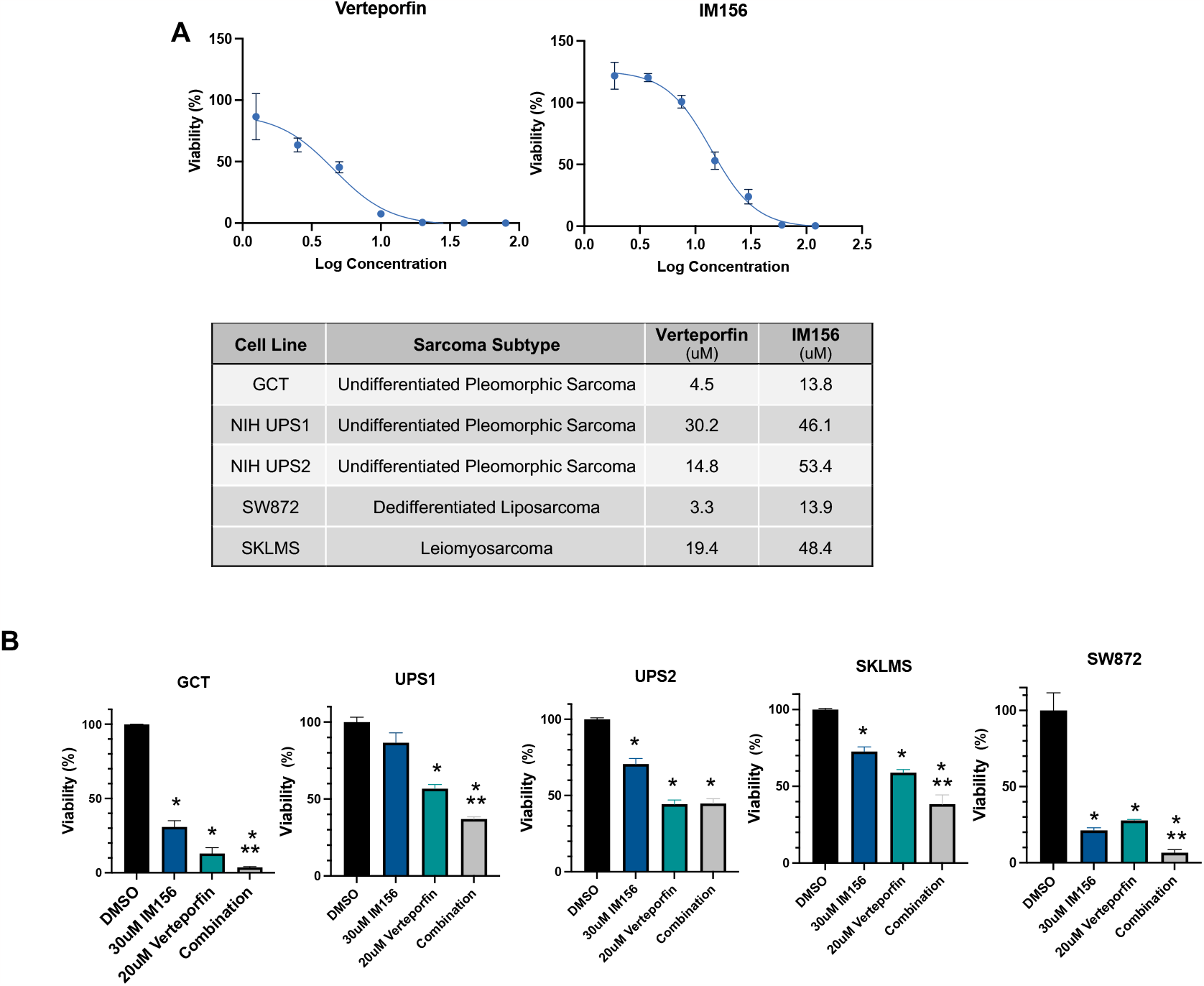
Treatment of Sarcoma Cell Lines with Hippo, Oxidative Phosphorylation or Dual Inhibition. (A) Top panel shows GCT (UPS cell line) proliferation with the addition of increasing concentrations of either the YAP1 inhibitor verteporfin or the complex I inhibitor IM156. Bottom panel provides relative IC50 values for each of the tested cell lines (also see supplemental figure 6). (B) Treatment of sarcoma cell line panel with 30uM of IM156, 20uM of verteporfin, or the combination. *p<0.05 as compared to the DMSO only control. **p<0.05 as compared to IM156 and verteporfin alone.

## Discussion

The process of tumorigenesis is complex and involves a series of genetic, epigenetic, and environmental factors to drive transformation^42^. We present a model of human sarcoma development that provides a basis to begin to unravel the pathophysiology of high-grade sarcoma development, growth and metastasis and provides a platform to identify new therapeutic targets.

### Modeling Human Sarcoma Development, Evolution, and Metastasis

Mesenchymal stem cells (MSCs), the presumed cell of origin for sarcomas have presented significant challenges when used as starting material for studies of human sarcomagenesis. Human MSCs have been historically difficult to transform presumably due to telomere maintenance and senescence^12^. In addition, there is variability across human MSC populations reflecting the heterogeneity of the human population. In our experience, we have observed significant differences between aliquots of commercially available MSCs in terms of proliferation rate and transduction efficiency leading to issues with reproducibility. To overcome these challenges, we utilized ASC52telo, an immortalized mesenchymal stem cell line as the starting material for transformation. While there are inherent limitations to the use of immortalized cell lines, this cell line allowed for reliable and reproducible results across a series of experiments. Interestingly, in our genomic analysis it was observed that this cell line has baseline aneuploidy which has been previously observed in human MSCs in culture ^39^. This raises the possibility that these cells are shifted closer to a neoplastic fate, due to specific acquired changes or a general tolerance of genomic instability than non-immortalized MSCs making them amendable to transformation. Nevertheless, these cells are in a pre-transformed state as defined by the inability to grow as a tumor in vivo. In the transcriptome analysis it was observed that these cells are distinct from the tumors that form and as a result, these cells provide a starting material with proper characteristics to study the critical steps of transformation.

From available literature, it is known that the process of sarcoma formation requires multiple hits^12^. The MSC-based forward genetics transformation model recreates this aspect of sarcoma biology. Both the loss of a transcription factor and the addition of an oncogenic gene were required for transformation. This provides the opportunity to study the cellular evolution of sarcoma development. In this study, twenty-seven factors were added but this can be scaled to include large classes of genes, specific pathways, or the whole genome. Despite the heterogeneity of primary sarcoma tumors, they uniformly metastasize to the lung which is the primary cause of mortality and this phenotype was observed in the model ^30^. We can use this system to address key clinical questions regarding the affinity of sarcoma to the lung, screen for drivers and regulators of metastasis and identify and test therapeutic targets.

### Identification of Drivers and Modifiers of Sarcoma Development

Using a genetic screen approach, several drivers of sarcoma formation were identified and then validated. *YAP1* and *KRAS* were clearly demonstrated to robustly transform MSCs into two common high-grade sarcomas, UPS and MFS. Additional drivers were identified including *PIK3CA* and *CDK4*. Efforts have been made to enhance the consistency of the *PIK3CA* and *CDK4* phenotypes, to generate a single histology from these drivers but this has not been achieved. The extended latency required for the growth of these tumors raises the question of spontaneous transformation which occurred after 6 months in a subset of implanted cells. While the *RB-/-P53+/-* cells were generated from a single cell clone, the loss of these tumor suppressors is likely increasing genomic instability leading to subpopulations with different genetic and epigenetic changes. We hypothesize that *CDK4* and *PI3KCA* provide a survival advantage to a subset of cells allowing spontaneous tumor formation while driving the development of an alternative histology. In previous studies we observed spontaneous tumors at a long latency that were UPS or OS histologically. Therefore, it is likely that *CDK4* can drive the formation of leiomyosarcoma but given the long latency and the survival advantage *CDK4* provides, a subpopulation of cells are able to spontaneously transform into UPS leading to two histologies from the same driver. Future studies using cell lineage tracing and spatial transcriptomics can delineate these two possibilities. This dual phenotype provides a unique opportunity to understand the juxtaposition of undifferentiated pleomorphic sarcoma with osteosarcoma and leiomyosarcoma. These data indicate a close relationship between these subtypes and that there are potentially subtle changes regulating cell fates that are being further investigated.

In the screen, there were two genes that were repeatedly identified in the barcode sequencing of the tumors but, did not give rise to tumors when transduced alone, *DDIT3* and *CCND1*. The addition of *DDIT3* to *KRAS* driven tumors resulted in a shorter latency and increased pleomorphism indicating a more aggressive tumor. These factors may function as modifiers of sarcoma development and promote tumor formation. Given the complex karyotype of sarcomas and the absence of single driver mutations, likely, a combination of genes contributes to the development of specific histologies and clinical behavior. Investigations into the roles of these modifier genes will provide additional insight into sarcoma biology and cellular pathways required for sarcomagenesis.

### Heterogeneity of Human Sarcomas is Reflected in the Model

The heterogeneity of sarcomas is multilayered. In addition to histology and patient related heterogeneity, intratumoral and intrasubtype heterogeneity also exist. This has been observed clinically with patients within one subtype such as undifferentiated pleomorphic sarcoma (UPS) having vastly different responses to the same therapies. It has been observed that sarcomas can contain more than one histology indicating they have undergone heterologous differentiation into another subtype. For example, a tumor that is primarily UPS but contains areas that are generating bone and appear histologically to be osteosarcoma^43^. This is not the case in the majority of patient tumors, but these cases are representative of the fact that sarcoma subtypes are likely a spectrum of disease with considerable plasticity^5^. In this model system, we see parallels to the heterogeneity that is seen across patients. While we observed UPS and MFS consistently in the *YAP1* and *KRAS* driven tumors, in those driven by *CDK4, PI3K*, and *JUN*, multiple or mixed histologies were seen. This is either driven by the heterogeneity of the starting material or to the evolution of the sarcomas over time which is being addressed in ongoing studies. In this study we have also observed that multiple genes can give rise to the same histology, UPS. Understanding the heterogeneity of those tumors and if this will provide further insight and overlap onto the human UPS disease spectrum warrants further investigation.

### Aneuploidy in Sarcoma Formation and Growth

Aneuploidy is the hallmark of non-single translocation driver sarcomas including the subtypes that are most common in adults such as UPS and MFS. The recapitulation of aneuploidy and copy number variations in general, and the involvement of specific patient sarcoma-associated amplification events (*YAP1*), are two key features of this model for the study of high-grade complex karyotype sarcomas. In cancer in general, aneuploidy has been implicated in key cellular processes such as tumorigenesis and metastasis^44^. In sarcoma, the impact of aneuploidy has not been extensively studied but will provide important insights into important clinical questions surrounding drug resistance and metastasis. We observed notable aneuploidy in the ASC52telo parental MSC line, and a further evolution of aneuploidy with transformation. Interestingly, it was observed that chromosome 11q which contains *YAP1* was amplified in tumors driven by *CDK4* and *PIK3CA* and this amplification was not detected in the cell lines used as starting material. These results support a tumor fitness benefit of *YAP1* amplification, with the amplification either occurring de novo or being selected for from a rare clone during the transformation process or during tumor growth. Based on the TCGA patient sarcoma data, there are a subset of human tumors that have amplified *YAP1*, and likely this is a key pathway driving the pathophysiology of a subset of high-grade sarcomas. *YAP1* is a transcription factor in the Hippo signaling pathway that has been shown to regulate mesenchymal stem cell and sarcoma proliferation, motility, and differentiation ^45^. Further investigation into the role of *YAP1* in driving sarcoma development and transformation may provide new mechanistic insights.

### Translational Potential of the Developed Model System

Within complex karyotype sarcomas, there are numerous genomic and transcriptomic changes and it is unclear which of these are driving the pathophysiology. The comparison of the tumors generated in the model to their patient counterparts allowed us to identify key pathways for sarcoma growth. We identified *YAP1* and oxidative phosphorylation as relevant clinical targets in a subset of high-grade sarcomas. Currently, YAP/TAZ and TEAD inhibitors are being tested in clinical trials and have shown promising early data in mesothelioma and tumors driven by Hippo pathway dysregulation^46^. In addition, IM156, the oxidative phosphorylation inhibitor has been tested in a phase I trial and led to stable disease in 7 out of 22 patients with limited side effects^47^. Based on the data from our model and the panel of high-grade sarcomas tested, these therapies could be effective for a subset of sarcoma patients and this warrants further investigation and consideration for early phase trials.

Through transcriptomic analysis comparing model tumors to patient tumors supported that YAP1- and KRAS-driven tumor lie near the endpoints of an unbiasedly defined spectrum of patient UPS and MFS tumors. The sarcoma community acknowledges that within each subtype and in particular, within UPS there are as yet unidentified subsets or continuous spectrums that could direct treatment decisions. It is feasible that the identified YAP1-like UPS-high subset may be highly sensitive to both inhibition of Hippo signaling and oxidative phosphorylation. Using this model, we can continue to expand our analysis and identify and define additional subgroups within high-grade sarcomas that may have unique vulnerabilities. While further perspectives and comparisons will need to be investigated, both unbiased and model-based transcriptome analyses stand to improve the current practice of classifying tumors in the undifferentiated MFS and UPS subtypes, and support a spectrum-based rather than binary-based approach.

Here, we have generated four sarcoma subtypes from a single cell of origin, undifferentiated pleomorphic sarcoma, myxofibrosarcoma, leiomyosarcoma, and osteosarcoma indicating that these subtypes have common underlying biology and may exist together on a disease spectrum. This model allows for additional insights into sarcoma biology and more broadly, can serve as a system for those that are tumor agnostic to understand critical aspects of cancer biology. The opportunities for further investigation and studies of high-grade complex karyotype sarcoma biology are vast and this preclinical model will serve as the basis to understand and target pathways and mechanisms driving the pathophysiology of this devastating disease.

## Acknowledgements

This work was supported in part by the UC Davis Paul Calabresi Career Development Award for Clinical Oncology as funded by the National Cancer Institute/National Institutes of Health grant #5K12-CA138464 and the Slifka Foundation. Research reported in this publication was supported by the National Cancer Institute of the National Institutes of Health under Award Number P30CA093373 (J.R.C.). Additional support was provided by The Doris Duke Charitable Foundation COVID-19 Fund to Retain Clinical Scientists awarded to UC Davis School of Medicine by the Burroughs Wellcome Fund (J.R.C.). Support was also provided by the NIH NCI RO1 CA222877, the UCLA SPORE in Prostate Cancer (NIH NCI P50 CA092131), the W.M. Keck Foundation, the UCLA Eli and Edythe Broad Center of Regenerative Medicine and Stem Cell Research Hal Gaba Director’s Fund for Cancer Stem Cell Research (T.G.G.). Research is supported by the Musculoskeletal Tumor Society Mentored Research Award (S.W.T.). Support was provided by the National Institutes of Health Grant #GM099134) and a Faculty Scholar grant from the Howard Hughes Medical Institute (K.P.). Research reported in this publication was supported by the National Institute of General Medical Sciences of the National Institutes of Health under Award Number T32GM145388 (J.F.). The content is solely the responsibility of the authors.

The authors thank Qian Chen and the UC Davis Center for Genomic Pathology Lab and the UCLA Translational Pathology Core Laboratory (TPCL) for histology and immunohistochemistry support. In addition, Jonathan Van Dyke and the UC Davis Flow Cytometry Shared Resource for flow cytometry and cell sorting supported by UC Davis CCSG (NCI P30CA093373). The authors thank the UC Berkeley QB3 Genomics (RRID:SCR_022170) core for next generation sequencing support.

pCW-Cas9 was a gift from Eric Lander & David Sabatini (Addgene plasmid # 50661 ; http://n2t.net/addgene:50661 ; RRID:Addgene_50661). LRG (Lenti_sgRNA_EFS_GFP) was a gift from Christopher Vakoc (Addgene plasmid # 65656 ; http://n2t.net/addgene:65656 ; RRID:Addgene_65656).

## Author Contributions

Conceptualization J.R.C., K.C., O.N.W., J.F., T.G.G. ; Methodology J.F., T.G.G., J.L., J.R.C.; Validation E.O., M.D.; Investigation M.M., J.F., E.O., S.T., T.G.G., J.R.C.; Writing – Original Draft, M.M., J.F., E.O., T.G.G., J.R.C.; Funding Acquisition O.N.W., T.G.G., J.R.C.; Visualization J.F., M.M.; Supervision L.R., R.C., K.P., K.C., O.N.W., T.G.G., J.R.C.

## Methods

### Cloning of CRISPR/Cas9 Constructs

Guide RNA sequences targeting RB1 (GCTCTGGGTCCTCCTCAGG), P53 (CCGGTTCATGCCGCCCATGC), or a control sequence (GTAATCCTAGCACTTTTAGG) were individually cloned into the LRG vector as previously described^24^. Sequences were previously used in a genome wide screen by Wang et al^22^. In order to combine these guides into one vector, the Gibson cloning method was used (Supplemental Figure 1C). The U6 promoter, guide sequence, and scaffold were amplified with custom primers (IDT) that contained a tag overlapping the LRG sequence flanking the EcoRI site. LRG vectors containing the control sequence or RB1 targeting sequence were cut with EcoRI and then ligated with the amplified sequence using the NEB HiFi Assembly kit (Cat#E5520). Constructs can be obtained at Addgene (in process).

### Lentivirus Production

To generate lentivirus, 293T cells (ATCC) were transfected with the 3^rd^ generation lentivirus packaging plasmids pMDL, pREV, and pVSVg^48^ and a lentivirus construct using calcium chloride transfection. Media containing virus was collected 48 hours after transfection, filtered through a 0.22μm filter and concentrated by centrifugation using Amicon 100kDa centrifugal filter units (Millipore Cat # UFC9100). Virus was titered using the Lenti-X p24 Rapid Titer Kit (Takara Cat #632200).

### Cell Lines

To generate the ASC52telo cell lines with *RB1* and *P53* gene knockout or controls, ASC52 cells were cultured in MSC basal media (Lonza Cat #PT-3001). ASC52 was validated using STR profiling. Lentivirus was generated from pCW-Cas9 (construct with doxycycline inducible Cas9) as described above and this was added to cells using polybrene and spinfection^22^. Cells were selected in puromycin and then single cells were seeded into 96 well plates. The clones were expanded and tested by western blot for Cas9 in the presence or absence of doxycycline. The clone with the tightest regulation of Cas9 was used for cell line generation. Lentivirus constructs containing CRISPR guides were transduced into cells using spinfection and polybrene. Cells were expanded and then sorted for green fluorescent protein (GFP) positivity. Individual GFP positive cells were sorted into 96 well plates to create single cell clones. Clones were expanded and tested by western blot to determine which had loss of *RB1* and/or *P53*. The DNA level mutation leading to the knockout was determined by PCR of the region containing the CRISPR target sequence and subsequent TOPO cloning and sanger sequencing (Invitrogen K2800J10). For drug studies, GCT, SKLMS, and SW872 were purchased from ATCC. NIHUPS1 (Lot 12654-015-R-J1-PDC) and NIH UPS2 (Lot 317291-083-R-J1-PDC) are patient derived tumor cultures provided by the NCI Patient Derived Models Repository (PDMR). These were all cultured in provider recommended medias and tested regularly for mycoplasma contamination.

### Flow Cytometry

Human mesenchymal stem cells (ATCC PCS-500-012) or ASC52telo cells (ATCC SCRC-4000) were suspended in 5%FBS/PBS solution and the following antibodies were added: APC-CD14 (BD 561708), APC-CD19 (BD 561742), PE-CD29 (Biolegend 303003), APC-CD34 (BD 560940), PE-CD44 (BD 555479), APC-CD45 (BD 560973), PE-CY7-CD73 (BD 561258), PERCP-CY5.5-CD90 (BD 561557), PE-CD166 (343903). Percentage positive was determined as compared to an unstained control using FloJo analysis of FACS plots.

### In Vitro Differentiation

To differentiate mesenchymal stem cells in vitro, the Lonza protocol was used. Briefly, adipogenic, osteogenic, and chondrogenic medias were prepared by addition of supplements (Cat #PT-3002, PT-3003, PT-3004). ASC52telo cells were plated at 1x10^4^ per cm^2^ for adipogenic differentiation and 1.5x10^3^ per cm^2^ for osteogenic differentiation. Cell numbers used were less than recommended by the manufacturer because these cells lack contact inhibition. For chondrogenic differentiation, cells were resuspended at a concentration of 1.6x10^7^ and 5ul was used per pellet. Differentiation medias were added and changed every three days for 14-21 days. Cells were fixed in 10% PBS buffered formalin then stained with oil red o, alizarian red, and alcian blue as previously described for visualization^49^.

### Western Blot

Cells were grown in adherent conditions and either left untreated or treated with 20uM doxorubicin for 24 hours. Cells were collected and lysed in 8M urea buffer containing protease inhibitors (Prometheus 18-427). Protein was quantified using a BCA assay (Pierce Rapid Gold BCA Protein Assay PIA53226). Blots were probed using antibodies against P53 (Cell Signaling 2524) or RB1 (Cell Signaling 9309) followed by Beta Actin (Invitrogen MA515739). Bands were visualized using chemiluniscence (Supersignal West Pico PLUS PI34577).

### Lentivirus Library Production

LentiORF clones were purchased from Sigma containing genes of interest in the TCR3 backbone (Millipore Sigma). All constructs are V5 tagged except for KRAS. Each construct was verified by western blot using a V5 antibody. Plasmids were maxiprepped, quantified, then pooled together for lentivirus production. Lentivirus was produced using the 3^rd^ generation system as described above. In brief, 293T cells were transfected, media containing virus was collected after 48 hours and filtered. The virus was concentrated by centrifugation and tittered for addition to cells.

### Primary Screen

All in vivo work was ethically performed and approved by the UC Davis Institutional Animal Care and Use Committee. Cells lines with *RB1* and *P53* targeted or a control line were transduced with the lentiviral library at MOI of 200 using spinfection and polybrene. Cells were expanded in culture for 3-5 days and then trypsinized, counted, and suspended in 60ul of a 1:1 mix of HBSS and Matrigel (Corning). These cells were injected subcutaneously into 6-8 week old immuocompromised NSG mice (Jackson Labs Cat#005557). Tumors were monitored by palpation and once tumors reached 1cm, frozen and formalin fixed samples were collected.

### Secondary Screen

To test the ability of individual oncogenes to drive tumor transformation, individual TRC3 constructs (Sigma) were transduced into cells using spinfection. There were grown in culture and expanded then trypsizined, counted, and suspended in 60ul of a 1:1 mix of HBSS and Matrigel then injected subcutaneously into NSG mice.

### Barcode Sequencing

For analysis of barcodes in tumor samples, DNA was extracted from paraffin embedded samples. Tumors were sectioned and stained by H&E to ensure the presence of viable tissue. The subsequent section was removed from the slide and DNA was extracted (Qiagen, Cat#56404). Barcodes were amplified using flanking primers (CTTGAAAGTATTTCGATTTCTTGGC and TCCAGAGGTTGATTGTCGAC) and KAPA HiFi HotStart ReadyMix (Cat#K2602) to form a 104bp product. Adapters for next generation sequencing were ligated to the PCR products using the KAPA Hyper Plus Kit and KAPA dual indexed adapter kit (Cat#K8727 and Cat#KK8514). Libraries were evaluated using a bioanalyzer and samples were submitted for next generation sequencing to UC Berkeley QB3 genomics and sequenced using Illumina technology. Barcodes were extracted from sequenced library fastQs by correcting for read direction using grep to sort, then taking the first 24bp of each read, and then determining the top 100 unique most frequent barcodes in each library using uniq in unix. Sequences were checked for matches within 1 levenshtein distance by building a dictionary of possible sequences in R software and then searching for matches to known transgene barcodes. The total of each transgene barcode detected per library was tabulated from this data and then visualized using the pheatmap library in R.

### Histology

Paraffin embedding, sectioning, and H&E staining was performed by the UC Davis Genomic Pathology Lab. SATB2, myogenin, and alpha SMA staining was performed by the UCLA TPCL facility. Vimentin staining was done according to the following protocol. Immunohistochemistry (IHC) staining was performed on formalin-fixed paraffin embedded (FFPE) tissue sections. Sections were de-paraffinized in Xylene, then re-hydrated. Antigen Unmasking Solution (Vector Laboratories, Cat# H-3300), citrate-based pH 6.0, was used for antigen retrieval. Endogenous peroxidase activity was blocked by incubating sections with BLOXALL (Vector, Cat# SP-6000) for 10 minutes. The M.O.M Mouse IgG Blocking Reagent from the M.O.M ImmPRESS Polymer Kit (Vector, Cat# MP-2400) was used to block endogenous staining. The Vimentin primary antibody (Agilent, Cat#M072529-2) was diluted 1:75 in 2.5% horse serum. Secondary antibody staining was done with the M.O.M ImmPress Horse Anti-Mouse IgG for 10 minutes. The ImmPACT DAB EqV (Vector, Cat# SK-4103) reagent was used for visualization. Sections were counterstained with hematoxylin and mounted with VectaMount AQ Mounting Medium (Vector, CAT# H-5501). All slides were interpreted by Dr. Morgan Darrow, a board-certified pathologist specializing in sarcomas.

### RNA Sequencing

Total RNA was isolated from human sarcoma models and controls using the RNeasy Mini Kit (Qiagen). Library preparation was performed by the Functional Genomics Laboratory (FGL), a QB3-Berkeley Core Research Facility at UC Berkeley. Total RNA samples were checked on Bioanalyzer (Agilent) for quality. Only high-quality RNA samples (RIN > 8) were used. At the FGL, Oligo (dT)_25_ magnetic beads (Thermofisher) were used to enrich mRNA. The treated RNAs were rechecked on Bioanalyzer for their integrity. The library preparation for sequencing was done on Biomek FX (Beckman) with the KAPA hyper prep kit for RNA (now Roche). Truncated universal stub adapters were used for ligation, and indexed primers were used during PCR amplification to complete the adapters and enrich the libraries for adapter-ligated fragments. Samples were checked for quality on an AATI (now Agilent) Fragment Analyzer. Samples were then transferred to the Vincent J. Coates Genomics Sequencing Laboratory (GSL), where quantitative PCR was used to calculate sequence-able molarity with the Kapa Biosystems Illumina Quant qPCR Kits. Libraries were pooled evenly by molarity and sequenced on an Illumina NovaSeq6000 150PE S4. Raw sequencing data were converted into fastq format, sample-specific files using the Illumina bcl2fastq2 software.

### Gene Expression PCA and PLSR Analyses

RNA-seq fastq files were processed through the TOIL pipeline (https://github.com/BD2KGenomics/toil-rnaseq)^50^ . TOIL processed and batch effect normalized gene expression data of TCGA patient tumors were acquired from the XenaBrowser (https://xenabrowser.net). For visualization, RSEM expected count data was upper quartile normalized and log2 transformed. Unsupervised principal component analysis (PCA) and supervised Partial Least Squares Regression (PLSR) were performed on protein-coding genes. PCA was performed centered and unscaled using the prcomp function in R. PLSR was performed unscaled using the pls function in R. Projections onto PCA/PLSR frameworks were done by multiplication of the projected sample expression profiles by the rotation matrix.

To systematically select a range of primary cancer types from the TCGA to be included in the pancancer gene expression comparisons, a PCA of only protein coding genes was performed with all 33 TCGA cancer types. A Euclidean distance was then calculated between the TCGA sarcoma cluster and every other cancer type to measure overall transcriptome likeness. The distance was calculated to the 11^th^ component, accounting for >50% of the variance. The distance contributed from each component was multiplied by that component’s percent variance explained. Cancer types were then ranked closest (skin cutaneous melanoma) to farthest (brain lower grade glioma) and every fifth was selected.

### Differential Expression Analyses

Differential expression analysis was performed on raw RSEM expected count data of protein-coding genes using DESeq2 (http://www.bioconductor.org/packages/release/bioc/html/DESeq2.html) ^51^. DESeq2 was used to compare MFS and UPS amongst patients, and then within the human transformation model. Ggplot2’s stat_density_2d was used to highlight regions of high and low overlap in the rank-rank plot. Rank rank hypergeometric overlap analysis was performed using the R package RRHO (https://www.bioconductor.org/packages/release/bioc/html/RRHO.html). The R package lattice was used to plot the hypergeometric p-values (https://cran.r-project.org/web/packages/lattice/index.html).

### Gene Set Enrichment Analysis (GSEA) and GSEA-Squared

Gene set enrichment analysis (GSEA) was performed using the fgsea package (https://github.com/ctlab/fgsea/). Gene ontology (GO), canonical pathways (CP), and hallmark (H) gene sets from MSgidDB were included. Genes were ranked by the signed and log2 transformed p-value calculated by DESeq2^52^. To investigate trends in related enriched or de-enriched gene sets, GSEA-Squared analysis was performed. After identifying broad pathways of interest from top Normalized Enrichment Score (NES)-ranked GSEA results (e.g., Oxidative Phosphorylation, DNA Damage), related key terms of interest were queried. All gene sets were ranked by NES and marked for if they contained a key term in the pathway name. To assess the distribution of a category of terms, KS tests were performed using ks.test.2 (https://github.com/franapoli/signed-ks-test)^53^. The keywords used are listed below:

| Category | Keywords |
| --- | --- |
| Oxidative Phosphorylation | OXIDATIVE PHOSPHORYLATION, ELECTRON TRANSPORT CHAIN, MITOCHONDRIAL COMPLEX, NADH DEHYDROGENASE, MITOCHONDRIAL LARGE RIBOSOMAL |
| DNA Damage | DNA REPAIR, DNA DAMAGE STIMULUS, FANCONI, END JOINING |

### Whole Exome Sequencing

gDNA was isolated from human sarcoma models and controls using the QIAamp DNA Mini Kit (Qiagen). Library preparation was performed by the Functional Genomics Laboratory (FGL), a QB3-Berkeley Core Research Facility at UC Berkeley. The DNA quality was checked on Fragment analyzer (Agilent) and fragmented to the desired length using Biouptor pico (Diagenode). The library preparation for sequencing was done on Biomek FX (Beckman) with the KAPA hyper prep kit for DNA (now Roche). Truncated universal stub adapters were used for ligation, and indexed primers were used during PCR amplification to complete the adapters and enrich the libraries for adapter-ligated fragments. Samples were checked for quality on an AATI (now Agilent) Fragment Analyzer. Samples were then transferred to the Vincent J. Coates Genomics Sequencing Laboratory (GSL), where quantitative PCR was used to calculate sequence-able molarity with the Kapa Biosystems Illumina Quant qPCR Kits. Libraries were pooled evenly by molarity and sequenced on an Illumina NovaSeq6000 150PE S4. Raw sequencing data were converted into fastq format, sample-specific files using the Illumina bcl2fastq2 software.^51^

### Copy Number Variation Analyses

Copy number variation (CNV) analysis of the human models was performed using CNVKit V0.9.9 (https://cnvkit.readthedocs.io/en/v0.9.9/) ^54^. CNVKit was run using an in-house pooled normal reference from normal samples sequenced under the same conditions at the Technology Center for Genomics & Bioinformatics (TCGB) at UCLA. Copy number segmentation data of TCGA samples were download from XenaBrowser (https://xenabrowser.net).

Integrated CNA score ^40^ for each sample was defined as:

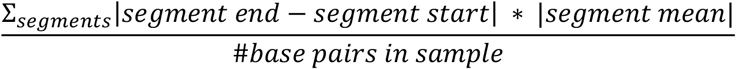

The copy number of a gene was calculated as follows with an assumed purity of 0.95 and ploidy of 2:

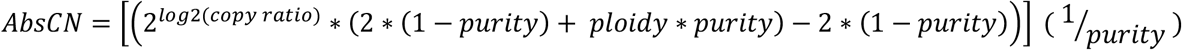

### Proliferation Analysis

For drug treatments, cells were plated into 96 cell plates on day 1 and treated with inhibitors or DMSO on day 2. The following day, relative proliferation was analyzed using Cell Titer Glo (Promega). Briefly, cells were lysed in the plate using the provided reagent and light was measured using a plate reader. Data was normalized to the DMSO control and plotted as a percentage of this value. IM156 (HY-136093A) and Verteporfin (HY-B0146) were both purchased from Med Chem Express.

