## Supplemental Figures 1-7 for "Genetic Screen in a Pre-Clinical Model of High-Grade Complex Karyotype Sarcoma Characterizes Drivers of Distinct Sarcoma Subtypes and Identifies New Therapeutic Vulnerabilities"

**Supplemental Figure 1.**

**A**

| Marker | Human MSC %Positive | ASC52telo % Positive |
|---|---|---|
| CD14 | 0 | 0 |
| CD19 | 0 | 0 |
| CD29 | 100 | 100 |
| CD34 | 0 | 0 |
| CD44 | 100 | 100 |
| CD45 | 0 | 0 |
| CD73 | 100 | 100 |
| CD90 | 100 | 100 |
| CD166 | 100 | 100 |

**B**

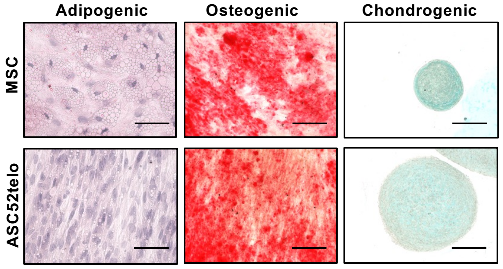


**C**

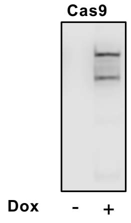


**D**

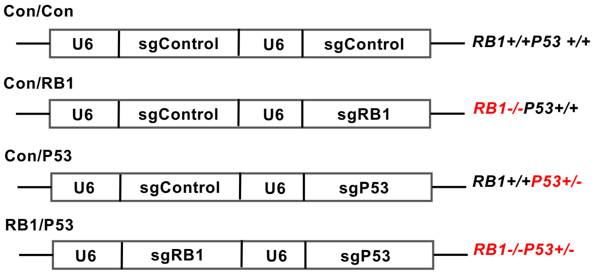


**E**

**RB1:** NM_000321, Target Sequence CTCCTGAGGAGGACCCAGAGC
**P53:** NM_000546, Target Sequence GCATGGGCGGCATGAACCGG

| Cell Line | *RB1* | *P53* |
|---|---|---|
| **Con/Con** *RB1+/+,P53+/+* | CTCCTGAGGAGGACCCAGAGC (Wildtype) | GCATGGGCGGCATGAACCGGAGG (Wildtype) |
| **Con/RB1** *RB1-/-,P53+/+* | CTCCT**T**GAGGAGGACCCAGAGC (c.249_250insT) CTCCT**TT**GAGGAGGACCCAGAGC (c.249_250insTT) | GCATGGGCGGCATGAACCGGAGG (Wildtype) |
| **Con/P53** *RB1+/+,P53+/-* | CTCCTGAGGAGGACCCAGAGC (Wildtype) | GCAT**T**GGGCGGCATGAACCGGAGG (c.870_871insT) GCATGGGCGGCATGAACCGGAGG (Wildtype) |
| **RB1/P53** *RB1-/-,P53+/-* | CTCCT**T**GAGGAGGACCCAGAGC (c.249_250insT) CTCCTGAGGAGGACCCAGAGC (c.249_250delGA) | GCAT**A**GGGCGGCATGAACCGGAGG (c.870_871insA) GCATGGGCGGCATGAACCGGAGG (Wildtype) |

**Supplemental Figure 1. Creation of Cell Lines Targeting *RB1* and *P53*** (A) Flow cytometry analysis of human mesenchymal stem cells and the immortalized ASC52 stem cell lines for markers of MSCs. Values shown are percentage of positive samples. (B) Histology of ASC52 cells after undergoing in vitro differentiation to adipocyctes, osteocytes, and chondrocytes as demonstrated by staining for Oil Red O, Alizarian red, and Alician blue respectively. Mesenchymal stem cells are shown in the top row as a positive control. Scale bar represents 300um (C) ASC52 cells were transduced with a construct containing doxycycline inducible Cas9. Western blot is shown for Cas9 expression in the absence and presence of doxycycline. (D) Schematic of cassettes created and cloned into the LRG vector to facilitate knockout of RB alone, P53 alone, or both. (E) Table showing verification of the CRISPR mediated knockout of each of the four cell lines.

**Supplemental Figure 2.**

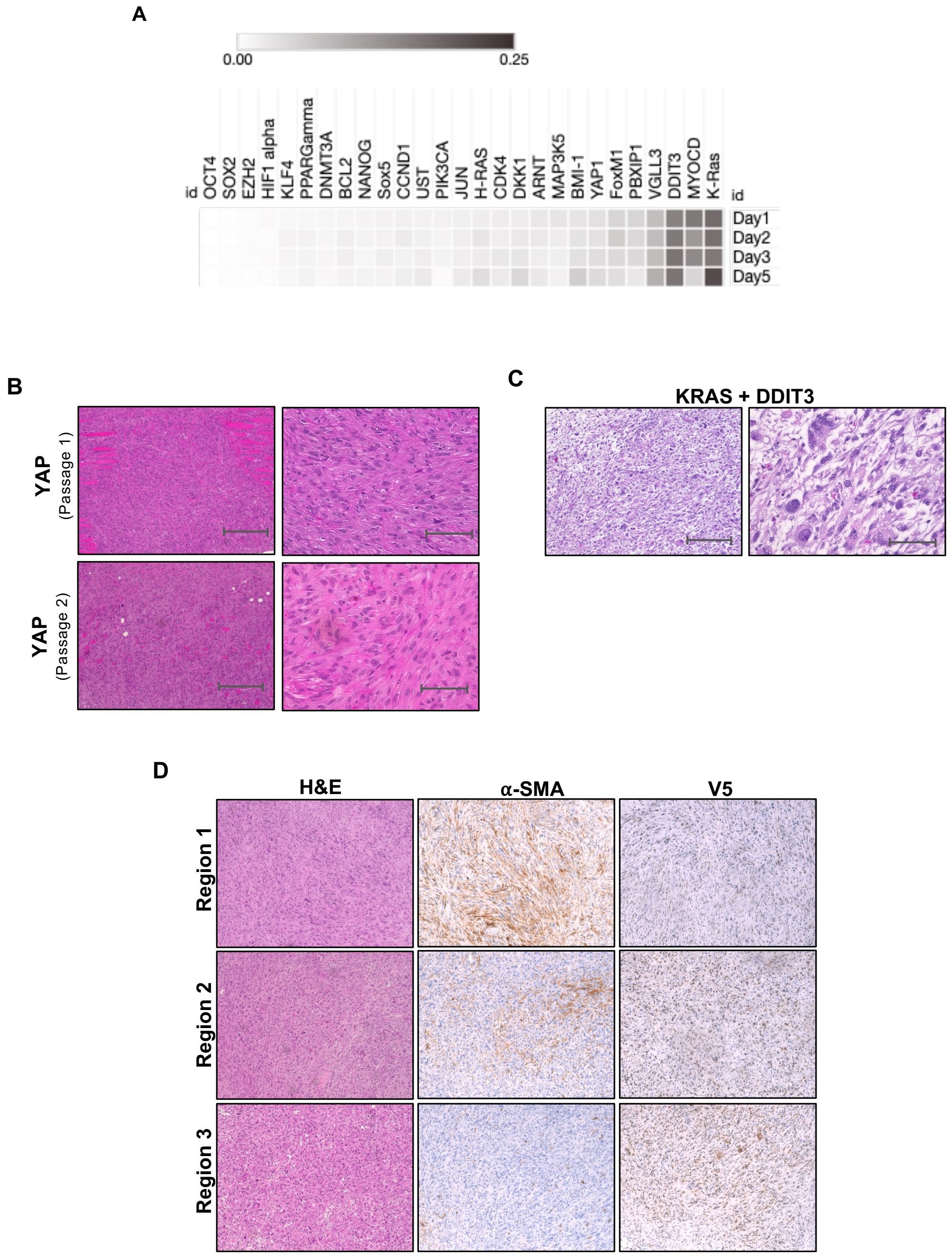


**Supplemental Figure 2. Primary and Secondary Screen Identify Drivers and Modifiers of Sarcoma Formation** (A) Heat map demonstrating the changes in expression of components of the lentiviral library over 5 days in culture. (B) H&E staining showing tumors expressing both KRAS and DDIT3 with increased pleomorphism. Low power (scale bar is 300um) is shown in the left column with high power in the right (scale bar is 50um). (C) Tumors generated from JUN demonstrating mixed histology by H&E staining. The presence of leiomyosarcoma can be seen with smooth muscle actin and V5 staining shows that the histology is not due to differences in JUN expression.

**Supplemental Figure 3.**

**A**

| | ARNT | BCL2 | BMI-1 | CCND1 | CDK4 | DDIT3 | DKK1 | DNMT3A | EZH2 | FoxM1 | H-RAS | HIF1a | JUN | K-Ras |
|---|---|---|---|---|---|---|---|---|---|---|---|---|---|---|
| **Starting 1** | 3.8 | 1.8 | 6.1 | 2.3 | 1.3 | 15.9 | 4.1 | 0.5 | 0.3 | 4.0 | 3.6 | 0.6 | 2.7 | 27.0 |
| **Starting 2** | 1.8 | 1.7 | 2.3 | 1.6 | 2.0 | 10.1 | 1.6 | 1.1 | 0.3 | 3.1 | 1.8 | 0.4 | 1.3 | 20.5 |
| **Starting 3** | 3.7 | 1.9 | 6.1 | 5.5 | 2.1 | 15.4 | 5.2 | 0.5 | 0.3 | 2.1 | 4.0 | 0.1 | 3.0 | 25.9 |
| **Tumor 1** | 0.0 | 0.0 | 0.0 | 0.0 | 0.0 | 8.3 | 0.0 | 0.0 | 0.0 | 0.1 | 0.0 | 0.0 | 0.0 | 3.5 |
| **Tumor 2** | 0.0 | 0.0 | 0.0 | 0.0 | 0.0 | 21.2 | 0.0 | 0.0 | 0.0 | 0.1 | 0.0 | 0.0 | 0.1 | 9.7 |
| **Tumor 3** | 0.0 | 0.0 | 0.0 | 0.0 | 0.0 | 18.2 | 0.0 | 0.0 | 0.0 | 0.1 | 0.0 | 0.0 | 0.0 | 8.5 |
| **Tumor 4** | 3.4 | 0.0 | 0.1 | 7.4 | 0.0 | 4.9 | 0.0 | 0.0 | 0.0 | 0.0 | 0.0 | 0.0 | 0.2 | 6.8 |
| **Tumor 5** | 0.5 | 0.0 | 0.0 | 0.0 | 0.4 | 1.1 | 11.1 | 0.0 | 0.0 | 0.0 | 0.0 | 0.0 | 13.4 | 41.4 |
| **Tumor 6** | 0.7 | 0.0 | 0.0 | 0.0 | 0.5 | 2.0 | 8.6 | 0.0 | 0.0 | 0.0 | 0.0 | 0.0 | 10.0 | 49.1 |
| **Tumor 7** | 1.1 | 0.0 | 0.0 | 0.0 | 0.8 | 2.8 | 9.7 | 0.0 | 0.0 | 0.0 | 0.0 | 0.0 | 13.1 | 39.0 |
| **Tumor 8** | 10.3 | 2.2 | 5.8 | 3.0 | 0.0 | 14.3 | 2.4 | 0.0 | 0.0 | 0.0 | 1.5 | 0.0 | 7.6 | 19.6 |
| **Tumor 9** | 5.9 | 0.8 | 3.1 | 6.5 | 0.2 | 17.5 | 5.7 | 0.2 | 1.2 | 0.6 | 4.2 | 0.0 | 11.8 | 19.0 |
| **Tumor 10** | 1.0 | 0.0 | 0.0 | 0.1 | 0.5 | 1.3 | 8.2 | 0.0 | 0.0 | 0.0 | 0.0 | 0.0 | 6.5 | 54.2 |
| **Tumor 11** | 1.6 | 0.0 | 0.0 | 0.0 | 1.0 | 3.2 | 7.7 | 0.0 | 0.0 | 0.0 | 0.0 | 0.0 | 7.1 | 50.1 |
| **Tumor 12** | 1.6 | 0.0 | 0.0 | 0.0 | 1.1 | 2.7 | 8.1 | 0.0 | 0.0 | 0.0 | 0.0 | 0.0 | 8.5 | 44.2 |
| **Tumor 13** | 3.4 | 0.1 | 0.0 | 5.5 | 0.1 | 4.9 | 0.0 | 0.0 | 0.0 | 0.3 | 0.1 | 0.0 | 0.1 | 35.4 |
| **Tumor 14** | 7.5 | 1.3 | 4.5 | 1.7 | 0.0 | 15.3 | 1.4 | 0.0 | 0.0 | 0.0 | 0.7 | 0.0 | 4.2 | 23.4 |
| **Tumor 15** | 4.7 | 0.5 | 2.1 | 7.6 | 0.2 | 21.6 | 5.2 | 0.1 | 0.8 | 0.7 | 2.7 | 0.0 | 6.8 | 23.8 |

| | KLF4 | MAP3K5 | MYOCD | NANOG | Oct4 | PBXIP1 | PIK3CA | PPARg | SOX2 | Sox5 | UST | VGLL3 | YAP1 |
|---|---|---|---|---|---|---|---|---|---|---|---|---|---|
| **Starting 1** | 0.8 | 0.1 | 0.1 | 1.9 | 0.0 | 1.9 | 0.2 | 0.9 | 0.2 | 1.9 | 3.3 | 8.2 | 6.4 |
| **Starting 2** | 1.4 | 2.0 | 9.2 | 1.2 | 0.0 | 3.4 | 4.4 | 4.5 | 4.6 | 4.7 | 4.8 | 5.0 | 5.1 |
| **Starting 3** | 0.8 | 0.9 | 0.0 | 1.5 | 0.1 | 0.7 | 1.6 | 1.8 | 0.3 | 0.9 | 2.0 | 5.6 | 7.8 |
| **Tumor 1** | 0.0 | 0.0 | 0.0 | 0.0 | 0.0 | 0.0 | 0.0 | 0.0 | 0.0 | 0.0 | 0.0 | 0.0 | 88.1 |
| **Tumor 2** | 0.0 | 0.0 | 0.0 | 0.0 | 0.0 | 0.0 | 0.0 | 0.0 | 0.0 | 0.0 | 0.0 | 0.0 | 68.9 |
| **Tumor 3** | 0.0 | 0.0 | 0.0 | 0.0 | 0.0 | 0.0 | 0.0 | 0.0 | 0.0 | 0.0 | 0.0 | 0.0 | 73.2 |
| **Tumor 4** | 0.1 | 0.0 | 0.0 | 0.0 | 0.0 | 0.0 | 0.0 | 0.0 | 0.0 | 0.1 | 35.1 | 0.3 | 41.5 |
| **Tumor 5** | 0.2 | 0.3 | 0.0 | 0.0 | 0.0 | 0.0 | 0.5 | 0.0 | 0.2 | 0.0 | 12.2 | 9.9 | 8.5 |
| **Tumor 6** | 0.0 | 0.9 | 0.0 | 0.0 | 0.0 | 0.0 | 0.5 | 0.0 | 0.3 | 0.0 | 10.8 | 9.9 | 6.5 |
| **Tumor 7** | 0.1 | 0.5 | 0.0 | 0.0 | 0.0 | 0.0 | 1.0 | 0.0 | 0.5 | 0.0 | 9.5 | 9.8 | 12.1 |
| **Tumor 8** | 0.2 | 0.0 | 0.0 | 0.0 | 0.0 | 0.0 | 2.3 | 0.0 | 0.0 | 25.3 | 3.1 | 0.0 | 2.4 |
| **Tumor 9** | 0.0 | 1.2 | 0.0 | 1.4 | 0.0 | 0.0 | 0.3 | 0.0 | 0.0 | 3.9 | 2.3 | 4.2 | 9.9 |
| **Tumor 10** | 0.1 | 0.7 | 0.0 | 0.0 | 0.0 | 0.0 | 2.1 | 0.1 | 0.3 | 0.0 | 8.7 | 8.8 | 7.5 |
| **Tumor 11** | 0.2 | 1.3 | 0.0 | 0.0 | 0.0 | 0.0 | 2.3 | 0.0 | 0.8 | 0.0 | 7.4 | 9.5 | 7.7 |
| **Tumor 12** | 0.2 | 0.7 | 0.0 | 0.0 | 0.0 | 0.0 | 1.5 | 0.0 | 1.0 | 0.0 | 9.1 | 8.7 | 12.4 |
| **Tumor 13** | 0.1 | 0.0 | 0.0 | 0.0 | 0.0 | 0.0 | 3.9 | 0.0 | 0.0 | 0.0 | 19.1 | 0.3 | 26.7 |
| **Tumor 14** | 0.0 | 0.0 | 0.0 | 0.0 | 0.0 | 0.0 | 1.6 | 0.0 | 0.0 | 35.2 | 1.5 | 0.0 | 1.6 |
| **Tumor 15** | 0.0 | 0.6 | 0.0 | 1.3 | 0.0 | 0.0 | 0.3 | 0.0 | 0.0 | 2.9 | 1.4 | 5.7 | 11.1 |

**B**

| | ARNT | BCL2 | BMI-1 | CCND1 | CDK4 | DDIT3 | DKK1 | DNMT3A | EZH2 | FoxM1 | H-RAS | HIF1a | JUN | K-Ras |
|---|---|---|---|---|---|---|---|---|---|---|---|---|---|---|
| **Tumor 1** | 1 | 1 | 1 | 1 | 1 | 1 | 1 | 1 | 1 | 1 | 1 | 9.90E-01 | 1 | 1 |
| **Tumor 2** | 1 | 1 | 1 | 1 | 1 | 7.21E-22 | 1 | 1 | 1 | 1 | 1 | 9.97E-01 | 1 | 1 |
| **Tumor 3** | 1 | 1 | 1 | 1 | 1 | 4.29E-07 | 1 | 1 | 1 | 1 | 1 | 9.99E-01 | 1 | 1 |
| **Tumor 4** | 1.02E-16 | 1 | 1 | 7.68E-141 | 1 | 1 | 1 | 1 | 1 | 1 | 1 | 1 | 1 | 1 |
| **Tumor 5** | 1 | 1 | 1 | 1 | 1 | 1 | 2.93E-283 | 1 | 1 | 1 | 1 | 1 | 1.00E-296 | 1.00E-296 |
| **Tumor 6** | 1 | 1 | 1 | 1 | 1 | 1 | 5.27E-175 | 1 | 1 | 1 | 1 | 1 | 4.53E-255 | 1.00E-296 |
| **Tumor 7** | 1 | 1 | 1 | 1 | 1 | 1 | 8.39E-205 | 1 | 1 | 1 | 1 | 1 | 1.00E-296 | 4.14E-211 |
| **Tumor 8** | 4.73E-194 | 1.01E-02 | 1.32E-41 | 5.43E-12 | 1 | 2.75E-20 | 5.83E-05 | 1 | 1 | 1 | 9.29E-01 | 1 | 7.53E-145 | 9.51E-01 |
| **Tumor 9** | 2.81E-61 | 1 | 2.30E-04 | 8.11E-92 | 1 | 1.32E-54 | 1.13E-65 | 1 | 1.14E-15 | 1 | 2.38E-26 | 1 | 1.80E-296 | 9.96E-01 |
| **Tumor 10** | 1 | 1 | 1 | 1 | 1 | 1 | 7.53E-20 | 1 | 1 | 1 | 1 | 1 | 8.63E-22 | 2.57E-208 |
| **Tumor 11** | 1 | 1 | 1 | 1 | 8.98E-01 | 1 | 1.39E-15 | 1 | 1 | 1 | 1 | 1 | 5.13E-26 | 5.82E-148 |
| **Tumor 12** | 1 | 1 | 1 | 1 | 7.95E-01 | 1 | 1.77E-18 | 1 | 1 | 1 | 1 | 1 | 4.51E-38 | 4.79E-86 |
| **Tumor 13** | 8.89E-01 | 1 | 1 | 8.29E-20 | 1 | 1 | 1 | 1 | 1 | 1 | 1 | 1 | 1 | 7.07E-27 |
| **Tumor 14** | 9.78E-18 | 9.78E-01 | 1 | 9.90E-01 | 1 | 7.96E-01 | 1 | 1 | 1 | 1 | 1 | 1 | 1.64E-05 | 1 |
| **Tumor 15** | 5.44E-03 | 1 | 1 | 4.84E-37 | 1 | 8.35E-16 | 3.27E-03 | 1 | 1.38E-04 | 1 | 9.98E-01 | 1 | 7.56E-24 | 1 |
| **Combined** | 2.54E-261 | 1 | 5.67E-28 | 8.01E-270 | 1 | 8.64E-94 | 6.16E-312 | 1 | 2.14E-07 | 1 | 2.08E-12 | 1 | 3.46E-312 | 3.81E-312 |

| | KLF4 | MAP3K5 | MYOCD | NANOG | OCT4 | PBXIP1 | PIK3CA | PPARg | SOX2 | Sox5 | UST | VGLL3 | YAP1 |
|---|---|---|---|---|---|---|---|---|---|---|---|---|---|
| **Tumor 1** | 1 | 1 | NaN | 1 | 9.87E-01 | 1 | 1 | 1 | 1 | 1 | 1 | 1 | 1.00E-296 |
| **Tumor 2** | 1 | 1 | NaN | 1 | 9.96E-01 | 1 | 1 | 1 | 1 | 1 | 1 | 1 | 1.00E-296 |
| **Tumor 3** | 1 | 1 | NaN | 1 | 9.98E-01 | 1 | 1 | 1 | 1 | 1 | 1 | 1 | 1.00E-296 |
| **Tumor 4** | 1 | 1 | 1 | 1 | 8.27E-01 | 1 | 1 | 1 | 1 | 1 | 1.00E-296 | 1 | 1.00E-296 |
| **Tumor 5** | 1 | 1 | 1 | 1 | 8.57E-01 | 1 | 1 | 1 | 1 | 1 | 4.67E-122 | 2.07E-62 | 4.11E-32 |
| **Tumor 6** | 1 | 1 | 1 | 1 | 8.04E-01 | 1 | 1 | 1 | 1 | 1 | 1.81E-76 | 5.14E-55 | 2.90E-07 |
| **Tumor 7** | 1 | 1 | 1 | 1 | 7.69E-01 | 1 | 1 | 1 | 1 | 1 | 2.55E-45 | 6.89E-49 | 2.61E-91 |
| **Tumor 8** | 1 | 1 | 1 | 1 | 7.12E-01 | 1 | 1 | 1 | 1 | 1.00E-296 | 1 | 1 | 1 |
| **Tumor 9** | 1 | 1 | 1 | 8.22E-02 | 7.08E-01 | 1 | 1 | 1 | 1 | 9.95E-01 | 1 | 9.93E-01 | 6.99E-42 |
| **Tumor 10** | 1 | 7.42E-07 | 1 | 1 | NaN | 1 | 1.23E-18 | 1 | 5.93E-02 | 1 | 1.69E-32 | 1.34E-01 | 1.06E-02 |
| **Tumor 11** | 1 | 4.04E-13 | 9.99E-01 | 1 | NaN | 1 | 7.27E-21 | 1 | 7.16E-06 | 1 | 2.59E-21 | 5.75E-03 | 4.55E-03 |
| **Tumor 12** | 1 | 5.89E-07 | 9.99E-01 | 1 | NaN | 1 | 9.50E-13 | 1 | 1.94E-08 | 1 | 7.69E-36 | 1.53E-01 | 1.19E-27 |
| **Tumor 13** | 1 | 9.99E-01 | 1 | 1 | NaN | 1 | 1.69E-38 | 1 | 1 | 1 | 3.15E-152 | 1 | 3.80E-190 |
| **Tumor 14** | 1 | 9.86E-01 | 9.99E-01 | 1 | NaN | 1 | 1.19E-13 | 1 | 1 | 1.00E-296 | 1 | 1 | 1 |
| **Tumor 15** | 1 | 1.18E-05 | 1 | 9.97E-01 | NaN | 1 | 3.27E-01 | 1 | 1 | 5.38E-04 | 1 | 1 | 7.56E-19 |
| **Combined** | 1 | 1.43E-15 | 1 | 1 | 1 | 1 | 9.58E-80 | 1 | 2.29E-04 | 8.12E-312 | 6.22E-312 | 1.73E-143 | 3.03E-312 |

*NaN indicates barcode was not present in starting material

**Supplemental Figure 3. Quantification of Primary Screen Barcodes.** (A) Table showing percent values of each barcode represented in tumor outgrowths as compared to the starting material. Heat map coloring is shown to demonstrate the areas of greatest enrichment. (B) Table of p-values for enrichment of each barcode relative to the starting values. P-values were calculated using a chi-squared test with continuity correction for equality of two proportions.

**Supplemental Figure 4.**

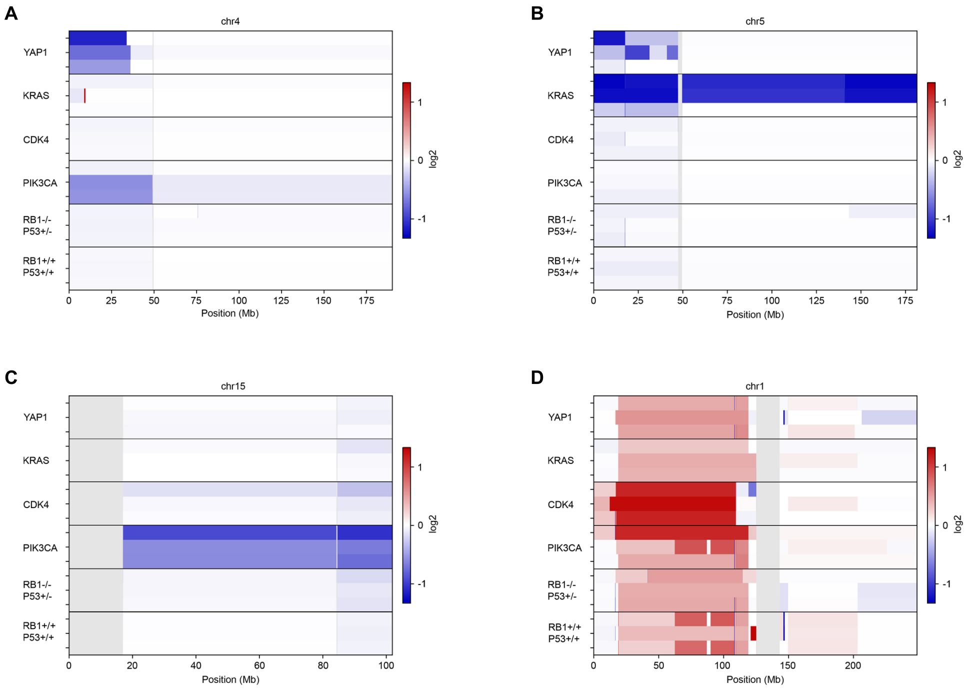


**Supplemental Figure 4. Chromosome Specific CNV.** Heat maps are shown of specific chromosome regions (A) Chromosome 4, (B) Chromosome 5, (C) Chromosome 15, (D) Chromosome 1 showing tumor specific chromosomal gains and losses as compared to the control cell lines

**Supplemental Figure 5.**

A

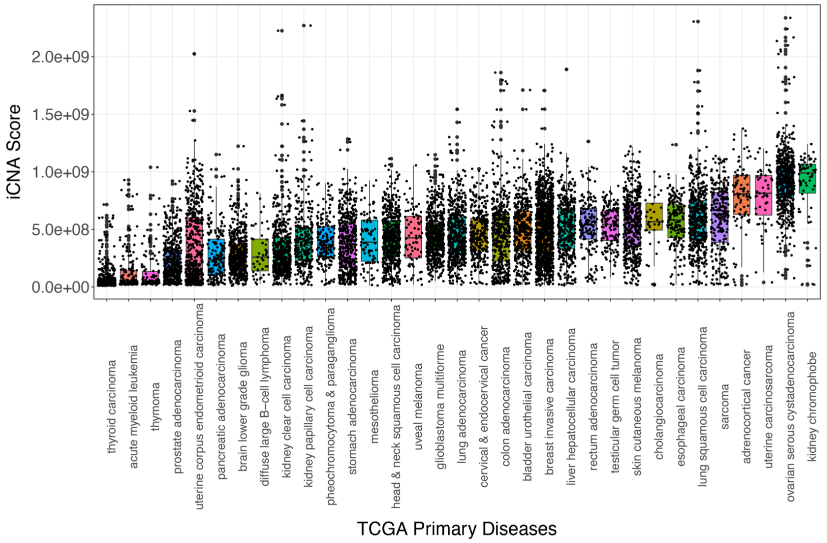


B

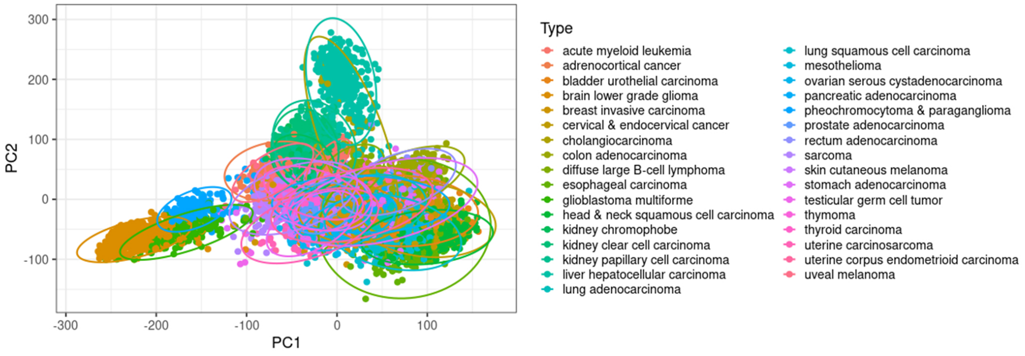


C

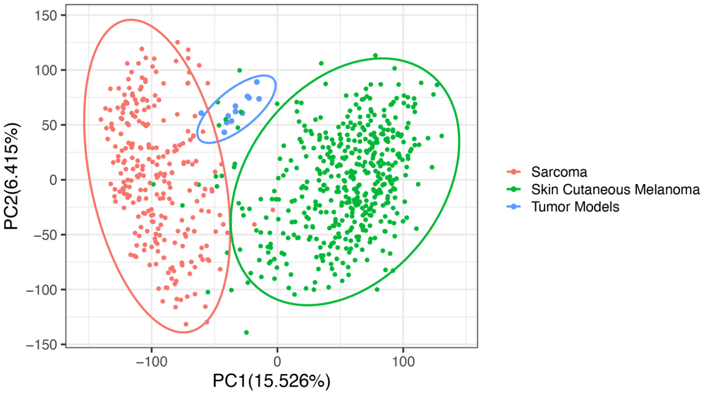


D

| Primary Disease | Distance |
|---|---|
| Sarcoma | 13.3652 |
| Skin Cutaneous Melanoma | 16.8146 |

**Supplemental Figure 5. Further Analysis of TCGA Data Including Additional Model and Patient Comparisons.** (A) iCNA score was calculated for all 33 tumor types sequenced by TCGA and plotted to show the high degree of aneuploidy seen in human sarcomas as compared to other tumor types. (B) PCA plot of the transcriptome of all TCGA analyzed tumors and the relative distance to sarcoma. (C) PCA plot of TCGA sarcoma and skin cutaneous melanoma gene expression data with projection of the tumor models onto the plot. (D) Euclidean distance calculation showing the tumor models project closer to sarcoma than skin cutaneous melanoma. (E) Heatmap of hypergeometric p-values generated from RRHO analysis.

**Supplemental Figure 6.**

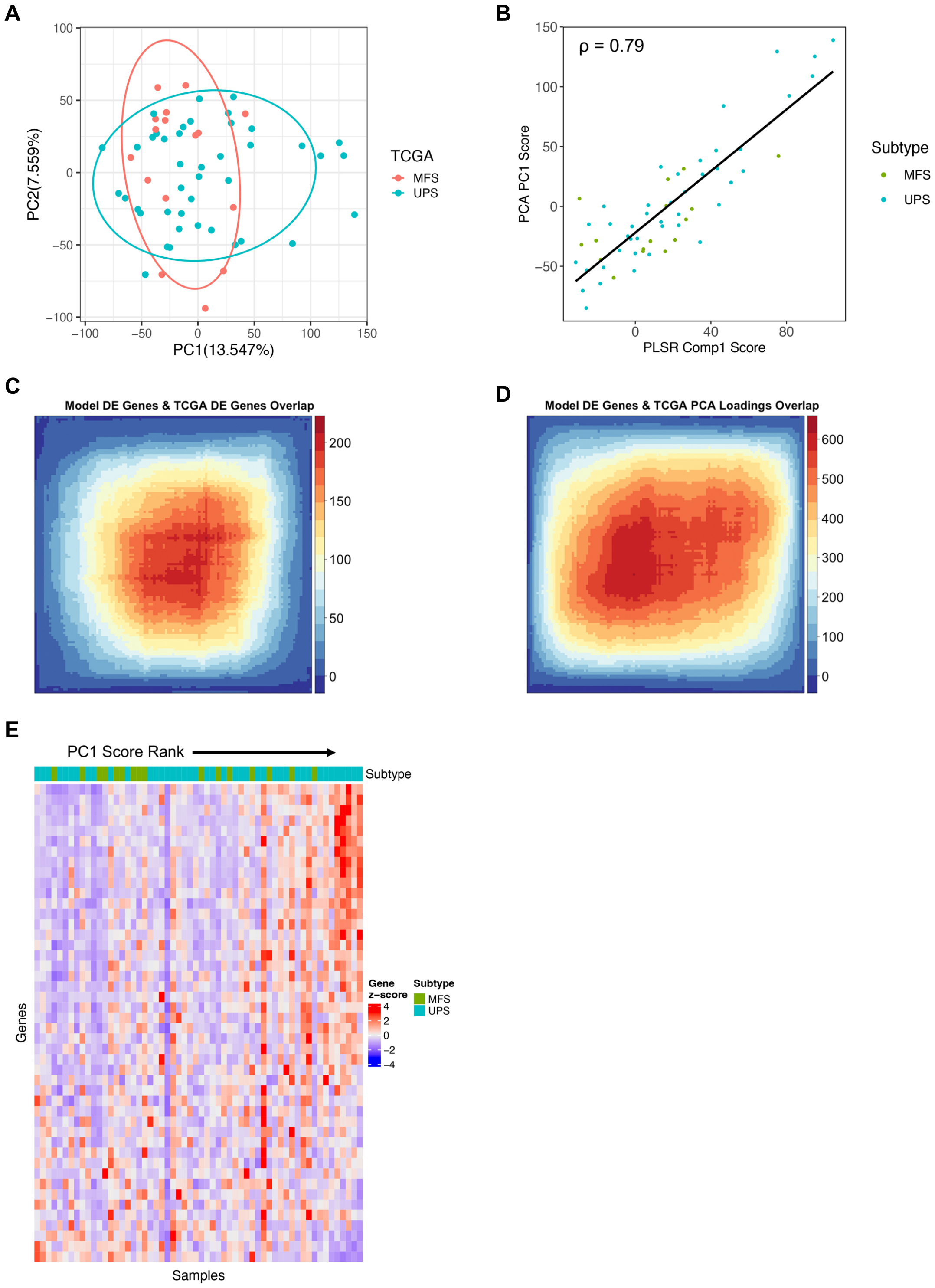


**Supplemental Figure 6. Further Analysis of Model Transcriptional Data.** (A) PCA plot of TCGA MFS and UPS samples. (B) Co-rank plot showing the correlation of MFS/UPS sample distribution between TCGA-defined PCA and model-defined PLSR. (C) Heatmap of hypergeometric p-values generated from RRHO analysis of model DE gene signature versus TCGA DE gene signature co-rank plot. (D) Heatmap of hypergeometric p-values generated from RRHO analysis of model DE gene signature versus TCGA PCA signature co-rank plot. (E) NDUF genes are positively enriched in UPS samples compared to MFS samples and in PCA loadings (+).

**Supplemental Figure 7.**

**A**

**Verteporfin**

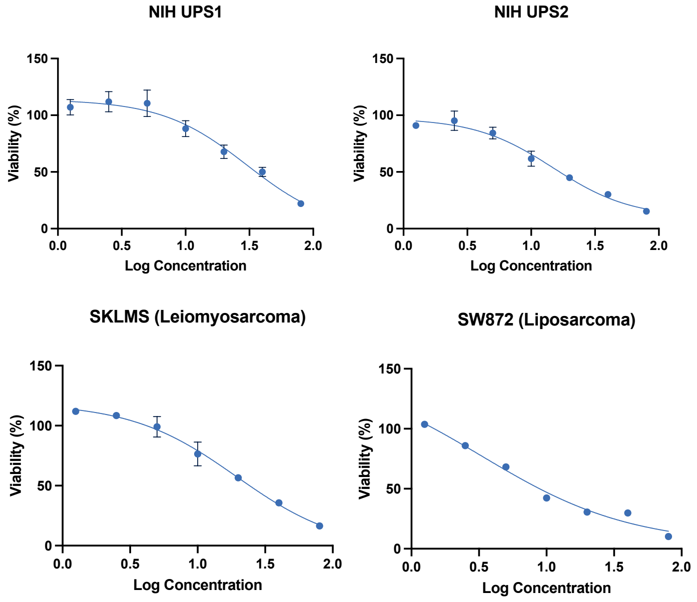


**B**

**IM156**

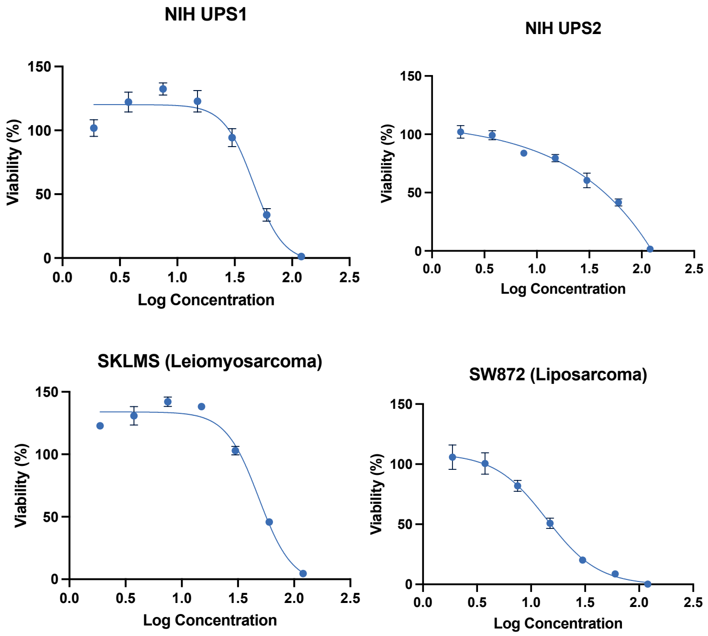


**Supplemental Figure 7. IC50 Curves.** A panel of cell lines were treated with increasing concentrations of (A) verteporfin or (B) IM156 for 24 hours and relative proliferation to the DMSO control were plotted. IC50 was determined by regression analysis using Prism.
